# Prevalence, persistence, and genetics of antibody responses to protein toxins and virulence factors

**DOI:** 10.1101/2021.10.01.462481

**Authors:** Julia W. Angkeow, Daniel R. Monaco, Athena Chen, Thiagarajan Venkataraman, Sahana Jayaraman, Cristian Valencia, Brandon M. Sie, Thomas Liechti, Payam Noroozi Farhadi, Gabriela Funez-dePagnier, Cheryl A. Sherman-Baust, May Q. Wong, Cynthia L. Sears, Patricia J. Simner, June L. Round, Priya Duggal, Uri Laserson, Theodore S. Steiner, Ranjan Sen, Thomas E. Lloyd, Mario Roederer, Andrew L. Mammen, Randy S. Longman, Lisa G. Rider, H. Benjamin Larman

**Affiliations:** Institute for Cell Engineering, Division of Immunology, Department of Pathology, Johns Hopkins University School of Medicine, Baltimore, MD, USA; Department of Biostatistics, Johns Hopkins Bloomberg School of Public Health, Baltimore, MD, USA; Department of Epidemiology, Johns Hopkins Bloomberg School of Public Health, Baltimore, MD, USA; ImmunoTechnology Section, Vaccine Research Center, NIAID, NIH, Bethesda, MD, USA; Environmental Autoimmunity Group, Clinical Research Branch, National Institute of Environmental Health Sciences, NIH, Bethesda, MD, USA; Jill Roberts Institute for Research in IBD, Division of Gastroenterology and Hepatology, Department of Medicine, Weill Cornell Medicine, New York, NY, USA; Laboratory of Molecular Biology and Immunology, NIH/National Institute on Aging, Baltimore, MD, USA; BC Children’s Hospital Research Institute, Vancouver, BC, Canada; Departments of Medicine and Oncology, Johns Hopkins University School of Medicine and Molecular Microbiology& Immunology, Bloomberg School of Public Health, Baltimore, MD, USA; Department of Pathology, Johns Hopkins University School of Medicine, Baltimore, MD, USA; Department of Pathology, Division of Microbiology and Immunology, University of Utah School of Medicine, Salt Lake City, UT, USA; Department of Genetics and Genomic Sciences, Icahn School of Medicine at Mount Sinai, New York, NY, USA; Department of Neurology, Solomon H. Snyder Department of Neuroscience, Johns Hopkins School of Medicine, Baltimore, MD, USA; Muscle Disease Unit, Laboratory of Muscle Stem Cells and Gene Regulations, National Institute of Arthritis and Musculoskeletal and Skin Diseases, NIH, Bethesda, MD, USA

## Abstract

Microbial exposures are crucial environmental factors that impact healthspan by sculpting the immune system and microbiota. Antibody profiling via programmable Phage ImmunoPrecipitation Sequencing (PhIP-Seq) provides a high-throughput, costeffective approach for multiplexed detection of exposure and response to thousands of microbial protein products. Here we designed and constructed a library of 95,601 56 amino acid peptide tiles spanning a subset of environmental proteins more likely to be associated with immune responses: those with “toxin” or “virulence factor” keyword annotations. PhIP-Seq was used to profile the circulating antibodies of ~1,000 individuals against this “ToxScan” library of 14,430 toxins and virulence factors from 1,312 genera of organisms. In addition to a detailed analysis of six commonly encountered human commensals and pathogens, we study the age-dependent stability of the ToxScan profile and use a genome-wide association study (GWAS) to find that the MHC-II locus modulates the selection of bacterial epitopes. We detect previously described anti-flagellin antibody responses in a Crohn’s disease cohort and identify a novel association between anti-flagellin antibodies and juvenile dermatomyositis (JDM). PhIP-Seq with the ToxScan library provides a new window into exposure and immune responses to environmental protein toxins and virulence factors, which can be used to study human health and disease at cohort scale.

## Introduction

Exposure to microbial protein products likely begins before birth, dramatically increases during the postnatal period, and then continues throughout life.^1,2^ The route, order, and degree of exposures to these environmental antigens, along with host genetics, nutrition, stress and the overall health status of the host, sculpt the ensuing adaptive immune responses or lack thereof.^3,4^ In turn, the immune responses to specific organisms may determine their capacity to colonize and/or cause disease in the host, the latter either via direct tissue damage or rather more complex, indirect mechanisms. To better understand environmental determinants of health and disease, it is thus critical to develop scalable tools to characterize immune responses to the microbial products encountered by humans throughout their lives.

An extraordinary effort has been underway to characterize the human microbiota and associate specific components with susceptibility to, or protection from, diseases that include atopic dermatitis, obesity, autoimmunity, neurodegeneration, cancer and response to immunomodulatory therapies.^5–7^ Most such studies rely on genome identification, via 16S or shotgun metagenomic sequencing. Compared to detection of immune responses, there are three key limitations to approaches that rely on the direct detection of nucleic acids. First, an infection may trigger a cascade of events and then resolve prior to future disease presentation, making it challenging to link the infection with the disease. Immune responses to an infection typically outlast the infections themselves, thus providing a temporal bridge to downstream consequences. Second, metagenomic studies are limited by sequencing depth and body site-specific sampling. In contrast, immune responses to low abundance microbes located anywhere on or in the body may be detectable via examination of circulating antibodies or immune cells. Finally, immune responses to an organism may elevate its likelihood of involvement in a disease process, versus organisms that are simply detected, their recognition by the host unknown. For these reasons, we propose that massively multiplexed profiling of immune responses to environmental protein antigens provides epidemiological information that is complementary to nucleic acid sequencing-based approaches. Phage ImmunoPrecipitation Sequencing (PhIP-Seq) is an antibody profiling technology that uses programmable bacteriophage (phage) display of synthetic oligonucleotides to encode peptides designed to tile across databases of proteins, and uses high-throughput DNA sequencing as the readout. PhIP-Seq has particularly favorable characteristics for epidemiological studies, including the size of the antigen database that can be represented (hundreds of thousands of peptides), the low per-sample cost of the assay (tens of dollars), and the sample throughput (hundreds of samples per automated run). Key limitations of PhIP-Seq, however, include the lack of highly conformational epitopes, the lack of post-translational modifications, and the finite complexity of the library.

We and others have used a pan-human viral library (VirScan) to characterize placental transfer of viral antibodies,^8^ link enteroviral infection with acute flaccid myelitis and type I diabetes,^9,10^ define SARS-CoV-2 epitopes,^11^ and in large studies of exposure and response to viruses in health and after HIV^12^ or measles infection.^13^ PhIP-Seq with a library spanning the proteins of the Allergome database (“AllerScan”) has recently been used to profile allergic IgE antibodies and their modulation in response to wheat oral immunotolerance therapy.^14^ While it would not be feasible to create a phage library that covers the millions of proteins expressed by the human microbiota,^15^ we have designed and constructed a novel library to represent a set of environmental proteins more likely to elicit immune responses: the 14,430 diverse proteins with “toxin” or “virulence factor” included as keywords in the UniProt database. This PhIP-Seq library of 95,601 56-amino acid peptides is therefore a unique representation of the environmental “toxome”/ “virulome”, which we refer to here as the “ToxScan’’ library.

We used protein A/G coated magnetic beads to immunocapture the phage-displayed peptides of the ToxScan library that react with serum antibodies from four cohorts composed of ~1,000 individuals in aggregate. Our results define a large number of previously undocumented, yet frequently recognized microbial epitopes. We describe the plasticity, yet relative stability of an individual’s ToxScan profile during their lifetime, and present evidence that selection of certain epitopes is dependent on host genetics – most notably in the Major Histocompatibility Complex class II (MHC-II) region of the Human Leukocyte Antigen (HLA) locus. Finally, in an effort to uncover microbial associations with disease, we detect previously described anti-flagellin antibodies in patients with Crohn’s disease, and report them for the first time in association with juvenile dermatomyositis (JDM). PhIP-Seq with the ToxScan library is thus an efficient new tool for linking environmental immunity with health and disease at cohort scale.

## Results

### Construction and characterization of the ToxScan Library

We used programmable T7 phage display to encode a library of all non-viral proteins included in the UniProt database with “toxin” (12,679 proteins representing clusters at >90% sequence identity) and “virulence factor” (all 2,133 unique proteins) keyword associations (downloaded November 2016). The resulting 14,430 unique protein sequences (436 archaeal, 7797 bacterial, and 5743 eukaryotic) were from 2,936 unique species of 1,312 genera. These sequences were used as input to the pepsyn library design software,^16^ which converted them into a set of 95,601 overlapping 56-mer peptide tiles, and reverse-translated them into corresponding DNA sequences. A library of encoding oligonucleotides was synthesized (Twist Bioscience) and cloned into the mid-copy T7 phage display system as described previously (**Fig. 1A**).^14,17,18^ The quality of the resulting ToxScan library was assessed via sequencing to a depth of 288-fold coverage. We detected 99.3% of the library peptides, and found them to be relatively uniformly represented (**Fig. 1B**). Serum antibody reactivity to the library was subsequently investigated using the standard protein A/G-based PhIP-Seq methodology.^16,17^

**Figure 1.**
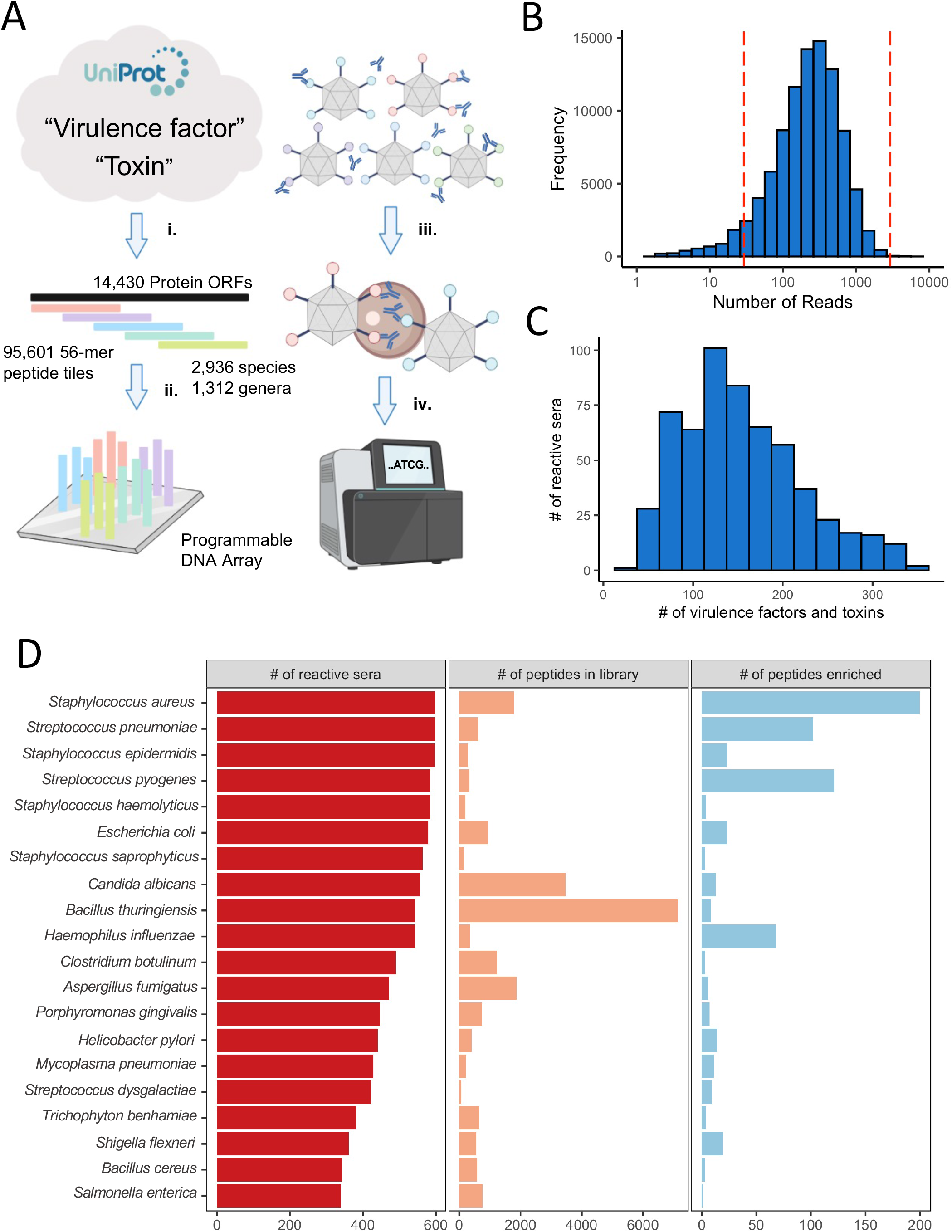
Antibody profiling with the T7 phage-displayed ToxScan library. **A. (i)** Protein sequences with “toxin” (collapsed to 90% sequence identity) or “virulence factor” (unique sequences) were downloaded from UniProt. **(ii)** The proteins were represented as 95,601 56 aa peptide tiles with 28 aa overlaps and encoded as 200-mer oligonucleotides. The library was cloned into the T7 phage display system and used for screening serum antibodies. **(iii)** IgG and bound phage are immunocaptured using magnetic beads. **(iv)** The library inserts are amplified by PCR and sequenced. **B.** ToxScan library QC: 99.3% of all library members were detected; 92.6% of the library was within one log (plus or minus) of the mean (indicated by vertical dashed lines). **C.** Number of ToxScan protein reactivities detected per individual in a healthy cohort (median=144, sd=66). **D.** Number of VRC samples reactive to at least 3 virulence factor or toxin peptides produced by an organism; the number of peptides from the organism in the ToxScan library, and the number of peptides enriched at a threshold of 5% prevalence (reactive in 30 or more individuals)

### ToxScan antibody profiles in a cross-sectional healthy cohort

We used the ToxScan library to profile plasma samples from a cross-sectional cohort of 598 study participants ranging in age from 18 to 70, which were collected by the National Institutes of Health Vaccine Research Center (VRC cohort). Basic demographic characteristics of these individuals are provided in **Table S1**. Across the cohort, at an individual level, we detected antibody responses to a median of 283 (standard deviation, sd = 118) peptides derived from a median of 144 (sd = 66) distinct proteins from 74 (sd = 36) species (**Fig. 1C**).

Among toxin and virulence factor antibody responses, we observed extensive immunoprevalence, yet diverse patterns of reactivity. The most frequently reactive library members were associated with prevalent human commensals and pathogens. Every individual in the VRC cohort had antibodies that reacted with at least three toxin or virulence factor peptides associated with *Staphylococcus aureus* (**Fig. 1D**). At a threshold of 5% seroprevalence (reactive in 30 or more individuals), 200 *S. aureus* peptides (of 1,775 in the library) from 53 proteins were considered immunoprevalent, exemplifying the detection of widespread, polyclonal responses to a variety of relevant epitopes. Notably, only 21% of these 200 reactive peptides shared sequence homology with linear antibody epitopes previously reported in the Immune Epitope Database (IEDB).^19^ All but two individuals had detectable responses to at least one *Streptococcus pneumoniae* peptide. 102 *S. pneumoniae* peptides (of 623 in the library) were considered immunoprevalent at the same threshold of 5% seroprevalence. We detected *Staphylococcus epidermidis* and *Streptococcus pyogenes* reactivities in 95% of the cohort. The most represented organism in the ToxScan library is *Bacillus thuringiensis*, an agriculturally important gram-positive bacterium found in soil that produces pesticidal crystal (Cry) proteins. Reactivities to *B. thuringiensis* toxins were frequently detected, but were found to be weak and lacked immunoprevalence. We observed immunoprevalent reactivity to an epitope in the heavy chain N-terminal translocation domain of *Clostridium tetani* toxin, presumably since most individuals had been immunized with either the DTaP or TdaP vaccines.

We used clustering to determine whether subsets of individuals could be grouped according to the similarity of their global reactivity towards prevalent epitopes. At a threshold of 5% seroprevalence, 848 peptides (0.89% of the 95,601 unique peptides assayed) were considered immunoprevalent (**Fig. 2A**). Two visually striking examples of minor clusters are annotated on the heatmap. Example cluster 1 is composed of *Staphylococcus* (both *S. aureus* and *S. epidermidis*) peptides derived from serine-aspartate repeat-containing proteins E, F, and G (SdrE, SdrF, and SdrG), and clumping factors A and B (ClfA and ClfB); these are functionally redundant surface proteins that share the ability to induce human platelet aggregation.^20^ IgG levels in sera for ClfB have been reported to be significantly higher in healthy individuals in comparison to acute-phase patients with clinically documented *S. aureus* infection, suggestive of protective immunity.^21^ A second example cluster is composed of related *Shigella flexneri* peptides, including the invasion plasmid antigens (Ipa) A, B, and C, which are associated with *Shigella* pathogenicity. These latter reactivities most likely reflect previous shigellosis in these specific individuals. In agreement with findings reported recently,^22^ *S. flexneri* seropositive individuals were significantly older than seronegative individuals (Mann-Whitney U Test, p=0.031). The highly seroprevalent peptides clustering at the bottom of **Fig. 2A** are largely composed of commonly encountered *Staphylococcus* and *Streptococcus* spp. toxins and virulence factors.

**Figure 2.**
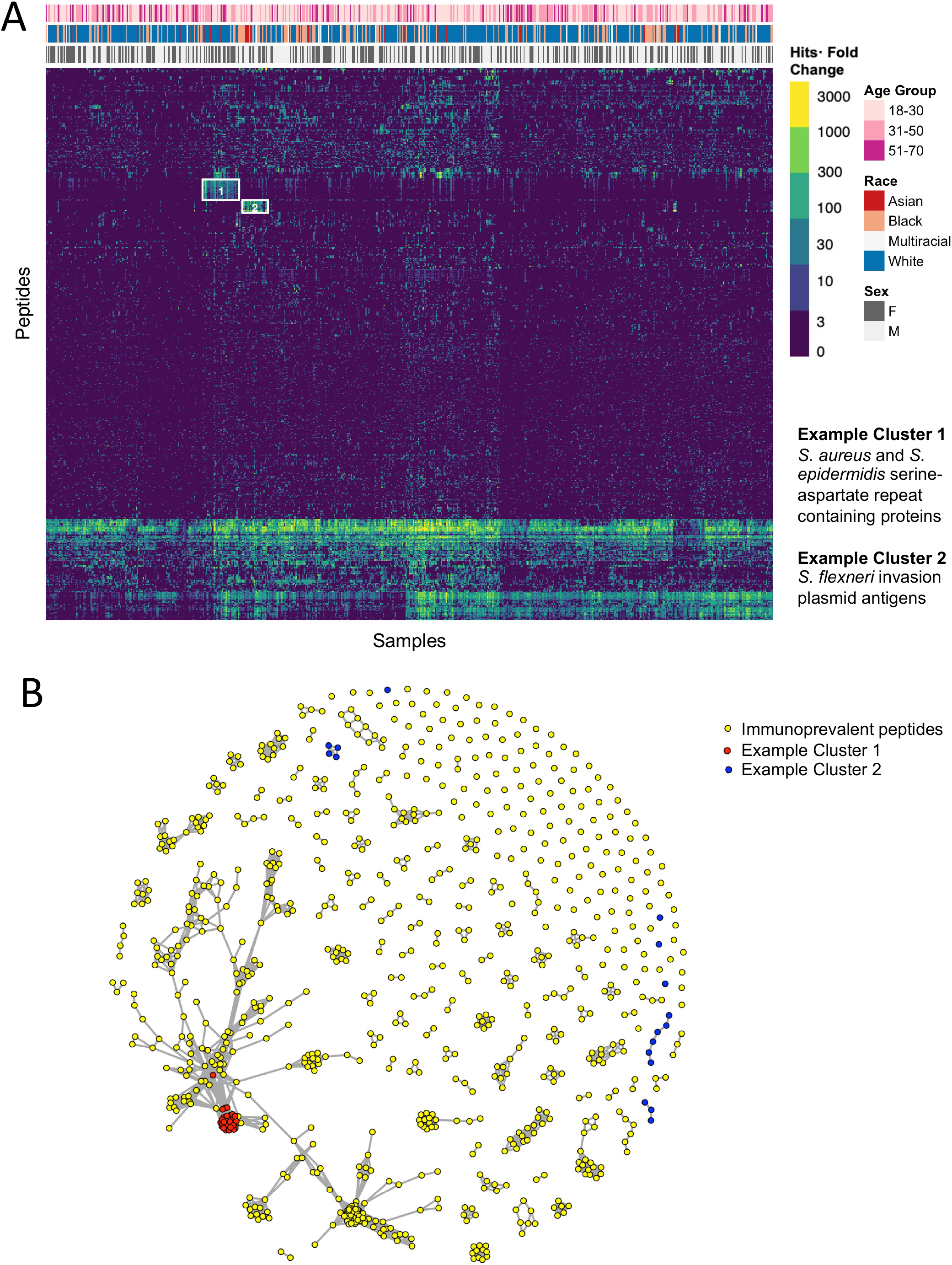
Immunoprevalent reactivities detected in a healthy human cohort. **A.** Heatmap showing the 848 immunoprevalent ToxScan peptide reactivities (at a threshold of 5% seroprevalence). Each row is a peptide and each column is a sample. Column annotations indicate the age, race, and sex of each individual. The color intensity of each cell indicates the fold change of each peptide versus mock immunoprecipitations (IPs). “Example cluster 1” is composed of *Staphylococcus aureus* and *S. epidermidis* serine-aspartate repeat-containing proteins E, F, and G (SdrE, SdrF, and SdrG), and clumping factors A and B (ClfA and ClfB). “Example cluster 2” is composed of *Shigella flexneri* peptides, including the invasion plasmid antigens A, B, and C (IpaA, IpaB, and IpaC). **B.** The 848 immunoprevalent peptides were aligned using blastp and a shared epitope network graph was constructed, with yellow dots corresponding to immunoprevalent peptides, red dots corresponding to “Example cluster 1” peptides and blue dots corresponding to “Example cluster 2” peptides.

In order to better understand homology shared between the observed immunoprevalent peptides and clusters, we used a network graph-based approach in which peptides (nodes) are linked (by an edge) if they share sequence similarity by blastp alignment (**Fig. 2B**). By calculating the maximal independent vertex set of the 848 immunoprevalent peptides, we determined that at least 364 distinct, public epitopes are represented among these peptides. Notably, 549 of the 848 immunoprevalent peptides share no sequence homology with any of the human host linear B cell epitopes previously reported in IEDB.

### Prevalent bacterial antibody responses

We next examined antibody recognition of toxins and virulence factors belonging to a subset of well-represented bacterial species in the library: *Streptococcus pneumoniae*, *Streptococcus pyogenes*, *Staphylococcus aureus, Staphylococcus epidermidis, Escherichia coli*, and *Haemophilus influenzae*. Only the immunoprevalent peptides are considered in the following analyses.

*S. pneumoniae* is a gram-positive commensal bacterium that is commonly found in the upper respiratory tract of humans, but which can also cause infections including pneumonia, bacteremia, otitis media, and meningitis.^23^ *S. pneumoniae* colonization is age-dependent; up to 27-65% of young children are carriers as compared to <10% of adults. We detected prevalent immunoreactivity targeting six *S. pneumoniae* virulence factors: immunoglobulin A1 protease (IgA1P), pneumococcal histidine triads D and E (PhtD and PhtE), pneumococcal surface protein A (PspA), and zinc metalloproteases B and C (ZmpB and ZmpC). We observed distinct subsets of individuals reactive to subgroups of IgA1P, PspA, and ZmpB epitopes. Strong and prevalent reactivity targeting PhtD and PhtE was observed, indicating that most healthy individuals are particularly reactive to these pneumococcal histidine triads **(Fig. 3A)**. A cluster of ZmpC and IgA1P peptides and several smaller sets of ZmpC and PhtE peptides have not been previously reported in IEDB (**Fig. 3B**).

**Figure 3.**
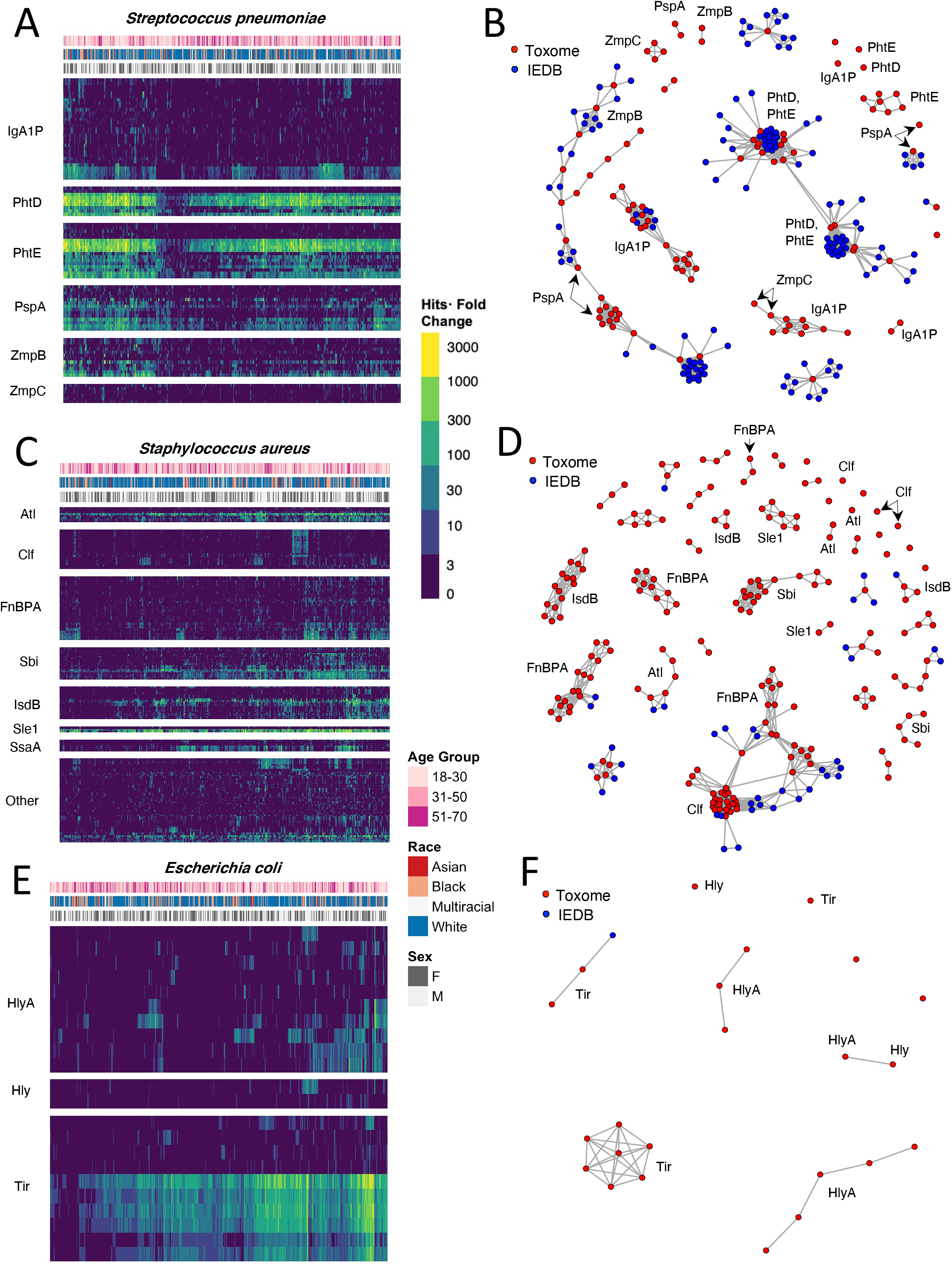
Immunoreactivity targeting proteins from three commonly encountered bacteria. Data in this figure are bacterial species-specific subsets of the VRC reactivities targeting the 848 immunoprevalent peptides. Column annotations and colors are as in **Fig. 2**. Column order is determined by hierarchical clustering. Row order is determined by hierarchical clustering, then subgrouped by protein family. **A.** Reactivities to *S. pneumoniae*: immunoglobulin A1 protease (IgA1P), pneumococcal histidine triads D and E (PhtD, PhtE), surface protein PspA, and zinc metalloproteases B and C (ZmpB, ZmpC). **B.** Shared epitope network graph depicting *S. pneumoniae* immunoprevalent ToxScan peptides and linear IEDB antibody epitopes. **C.** Reactivities to *S. aureus:* bifunctional autolysin (Atl), clumping factor (Clf), fibronectin binding protein A (FnBPA), immunoglobulin-binding protein Sbi, iron-regulated surface determinant B (IsdB) protein, N-acetylmuramoyl-L-alanine amidase (Sle1), and Staphylococcal secretory antigen (SsaA). **D.** Shared epitope network graph depicting *S. aureus* immunoprevalent ToxScan peptides and linear IEDB antibody epitopes. **E.** Reactivities to *E. coli:* alphahemolysin (HlyA), hemolysin (Hly), and translocated intimin receptor (Tir). **F.** Shared epitope network graph depicting *E. coli* immunoprevalent ToxScan peptides and linear IEDB antibody epitopes.

*S. pyogenes* (Group A *Streptococcus*) is a gram-positive extracellular pathogen that can colonize the skin and mucous membranes, particularly in the throat. *S. pyogenes* infections are the cause of many important infections and infectious complications, including bacterial pharyngitis (“strep throat”), scarlet fever, rheumatic fever, glomerulonephritis, streptococcal toxic shock syndrome, type II necrotizing fasciitis, cellulitis, erysipelas, and impetigo.^24^ Individuals of the VRC cohort displayed prevalent IgG reactivity to four virulence factors produced by *S. pyogenes*: C5a peptidase (SCPA), M proteins (M12, M2.1, M2.2, M24, M49, M5 and M6), streptolysin O (SLO), and virulence-factor related M protein (VrfM). We noted particular subgroups of individuals with increased reactivity to SCPA and M5 (**Fig. S1A**). Clusters of overlapping SCPA peptides have not been previously reported in IEDB **(Fig. S1B)**.

A gram-positive commensal bacterium, *S. aureus* typically colonizes the skin and mucosal surfaces, including the oropharynx. *S. aureus* is the leading cause of bacteremia and bone, joint, and surgical site infections, and can lead to important complications such as toxic shock syndrome; moreover, methicillin-resistant strains (MRSA) have emerged to cause significant morbidity and mortality.^25^ We detected reactivity to many *S. aureus* virulence factors and toxins, most notably bifunctional autolysin (Atl), clumping factor (Clf), fibronectin binding protein A (FnBPA), immunoglobulin-binding protein Sbi (Sbi), iron-regulated surface determinant B protein (IsdB), N-acetylmuramoyl-L-alanine amidase sle1 (Sle1), and staphylococcal secretory antigen A (SsaA, **Fig. 3C**). We observed near universal recognition of epitopes located in Sle1 and Alt. Sequence-based homology analysis demonstrated multiple clusters of FnBPA, Sbi, IsdB, and Sle1 epitopes not previously reported in IEDB (**Fig. 3D**). *S. epidermidis* and *S. aureus* can produce many of the same virulence factors but *S. epidermidis* lacks the enzyme coagulase and is comparatively less virulent.^26^ At a threshold of 5% seroprevalence, 23 of the 239 *S. epidermidis* peptides in the library (9.6%) were immunoprevalent, while 200 of the 1775 *S. aureus* peptides in the library (11.2%) were immunoprevalent (**Fig. 1D**). Similar to *S. aureus* Sle1, *S. epidermidis* Sle1 was universally recognized in the cohort, while SsaA was recognized in a subset of individuals (**Fig. S1C-D**).

*E. coli* is a gram-negative bacterium frequently found as a commensal in the human gastrointestinal tract, but many virulent pathotypes cause diarrheal illness, urinary tract infections, and invasive infections such as bacteremia and neonatal meningitis. For example, *E. coli* serotype O157:H7 and other Shiga toxin-producing *E. coli* (STEC) can cause hemorrhagic colitis and hemolytic uremic syndrome.^27,28^ Along with enteropathogenic *E. coli* (EPEC), which causes infant diarrhea, these enterohemorrhagic strains (EHEC) utilize a type III secretion system to inject the virulence factor, translocated intimin receptor (Tir), into the host cell. We detected near universal reactivity to ten Tir peptides (three non-homologous sets), and less reactivity against alpha-hemolysin (HlyA) and hemolysin (Hly) (**Fig. 3E**). Only one of these immunoprevalent peptides have been previously reported in IEDB (**Fig. 3F**).

*H. influenzae* is a gram-negative bacterium that can be pathogenic in humans. *H. influenzae* is categorized as either nontypeable or typeable (more likely to become invasive) according to the absence or presence of a polysaccharide capsule, respectively. In our library, the toxins and virulence factors produced by *H. influenzae* correspond to those that are conserved among all serotypes regardless of encapsulation. We detected prevalent, yet distributed reactivity to immunoglobulin A1 protease (IgAP), and less reactivity to outer membrane protein (OMP) (**Fig. S1E**). The numbers of seroprevalent clusters and singleton peptides in ToxScan analysis exceeded those in IEDB (**Fig. S1F**).

### Longitudinal stabilities of ToxScan profiles

We assessed factors contributing to the overall diversity of ToxScan peptide reactivities in the VRC cohort plasma. In this analysis, diversity was defined as the absolute number of ToxScan reactivities, regardless of overlap or prevalence. We considered sex, age, race and CMV status. On average, men had 10.82 more peptide reactivities than females (one-way ANCOVA, p = 0.02) after accounting for the other covariates. Neither race nor CMV status were significantly associated with the number of reactive peptides (p = 0.77, n = 152 for Black; p = 0.31, n = 23 for Asian; p = 0.42, n = 21 for multiracial; p = 0.23 for CMV status, 49.3% seropositive). Age, however, was independently associated with a decline of 0.89 reactivities per year (one-way ANCOVA, p = 2.8×10^-6^, **Fig. 4A**).

**Figure 4.**
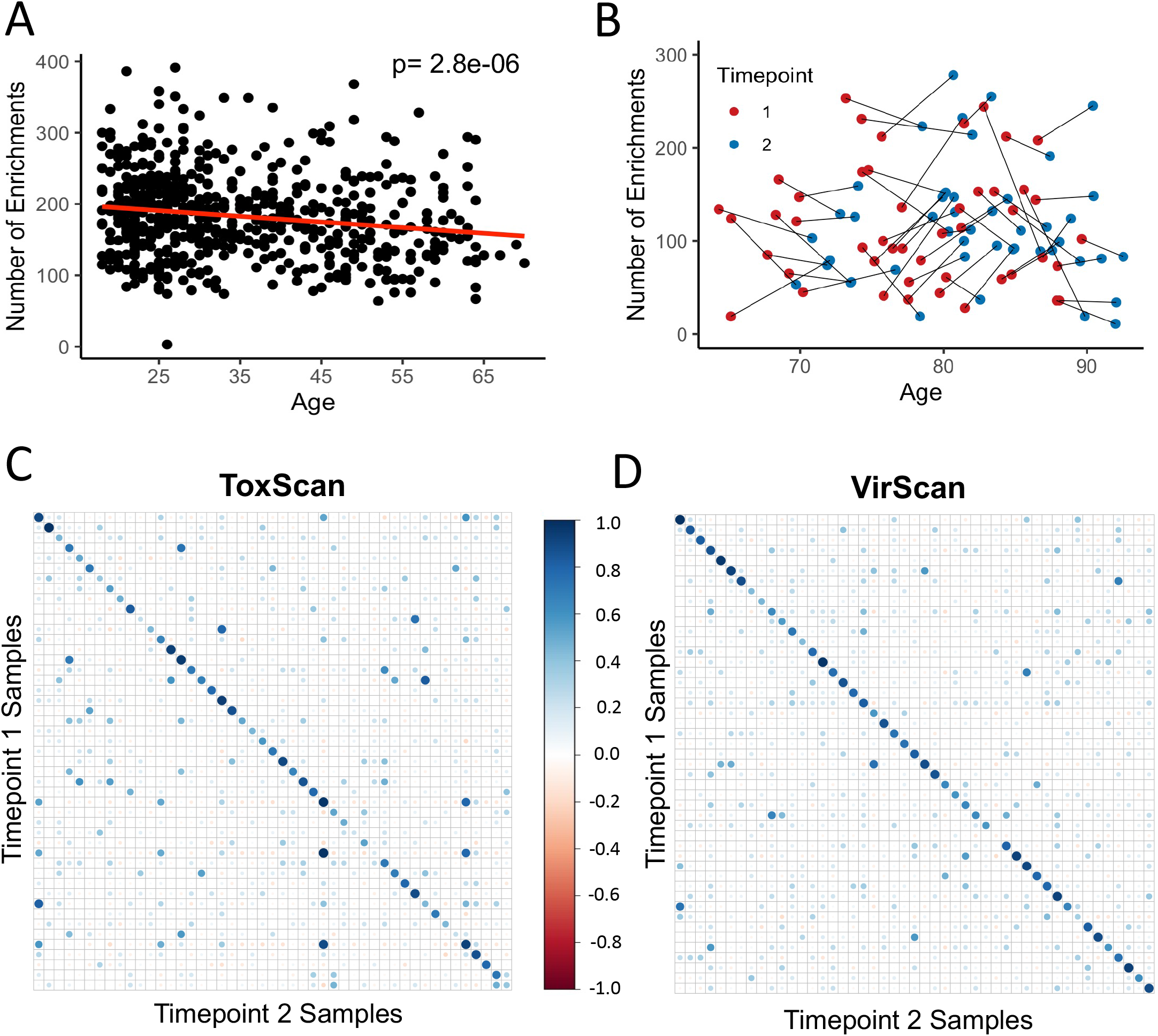
Longitudinal stability of ToxScan reactivities. **A.** Total number of reactive ToxScan peptides versus age (VRC cohort) reveals a decline of 0.89 reactivities per year (one-way ANCOVA, p= 2.8×10^-6^) after adjusting for covariates (sex, age, race, CMV status). **B.** The diversity of ToxScan reactivities over ~5 years in older adults (BLSA cohort). The first time point is indicated in red, the second in blue. ToxScan (**C**) and VirScan (**D**) longitudinal correlations of public peptide reactivities over ~5 years in older adults (BLSA cohort).

In a separate, longitudinal cohort composed of 47 healthy older volunteers aged ~70-90 from the Baltimore Longitudinal Study of Aging (BLSA, **Table S2**), we compared ToxScan profiles of individuals sampled at two timepoints roughly 5 years apart. The diversity of reactive ToxScan peptides was highly correlated (**Fig. 4B**, R=0.6, p=3.8×10^-6^), suggesting that humans tend to acquire an individual-specific ToxScan reactivity diversity set point, which is relatively stable, at least on the scale of several years in older adulthood.

In order to assess the intra-versus inter-individual patterns of immunoprevalent ToxScan reactivities, all longitudinal pairs of samples were compared (**Fig. 4C**). The same analysis was performed for immunoprevalent VirScan reactivities (**Fig. 4D**). Intraindividual sample pairs exhibited much higher levels of correlation versus interindividual samples pairs, for both ToxScan and VirScan. These data indicate that in addition to the ToxScan diversity set point, patterns of ToxScan reactivity are personal and persist over years. Both intra-individual and inter-individual sample pairs exhibited significantly higher levels of correlation among VirScan profiles versus ToxScan profiles (t-test, p= 0.038 and p=0.0012, respectively), potentially due to greater antigenic diversity among the seroprevalent peptides in the ToxScan library.

### Genome-wide association study of ToxScan peptide reactivity

We next set out to determine whether host genetics could influence the selection of epitopes within the ToxScan library. To this end, 7,637,921 variants were tested for association with any of the 166 ToxScan reactivities having immunoprevalence between 16% and 80% of the population (**Fig. S2**). The majority (132 of 166) of immunoprevalent reactivities targeted *S. aureus* (n=68) and *S. pyogenes* (n=64). Single-variant genomewide association studies (GWAS) were performed on each peptide, and the p-value threshold was adjusted for multiple hypothesis testing (Methods). The 478 individuals of European (EUR) descent in the VRC cohort were used for discovery and the 147 individuals of African ancestry (AFR) in the VRC cohort were used for replication (**Fig. S2**). Three peptides were associated with the class II Major Histocompatibility Complex (MHC-II) of the Human Leukocyte Antigen (HLA) locus on chromosomes 6, and one peptide was associated with the nucleoredoxin gene (*NXN*) on chromosome 17 (**Fig. 5A**). All three MHC-II associated peptides are from *S. aureus*; two adjacent, overlapping peptides are derived from the clumping factor A (ClfA) protein and the third is from Staphylococcal secretory antigen A1 (ssaA1). The variants associated with these three peptides replicated in the AFR sub-cohort. A credible sets analysis linked *HLA-DQA1* and *HLA-DQB1* to these *S. aureus* reactivities (**Fig. 5B-C, Fig. S3, Table S3**). The *NXN* associated reactivity to *S. pyogenes* M3 Streptolysin O (SLO) protein (p-value = 2.4×10^-10^ in EUR sub-cohort) could not be independently tested for replication, due to a skewed allele frequency in the AFR sub-cohort (0.986). These findings support the hypothesis that heritable factors, including those in the MHC-II locus, contribute to the specificity of human antibodies directed at microbial toxins and virulence factors.

**Figure 5.**
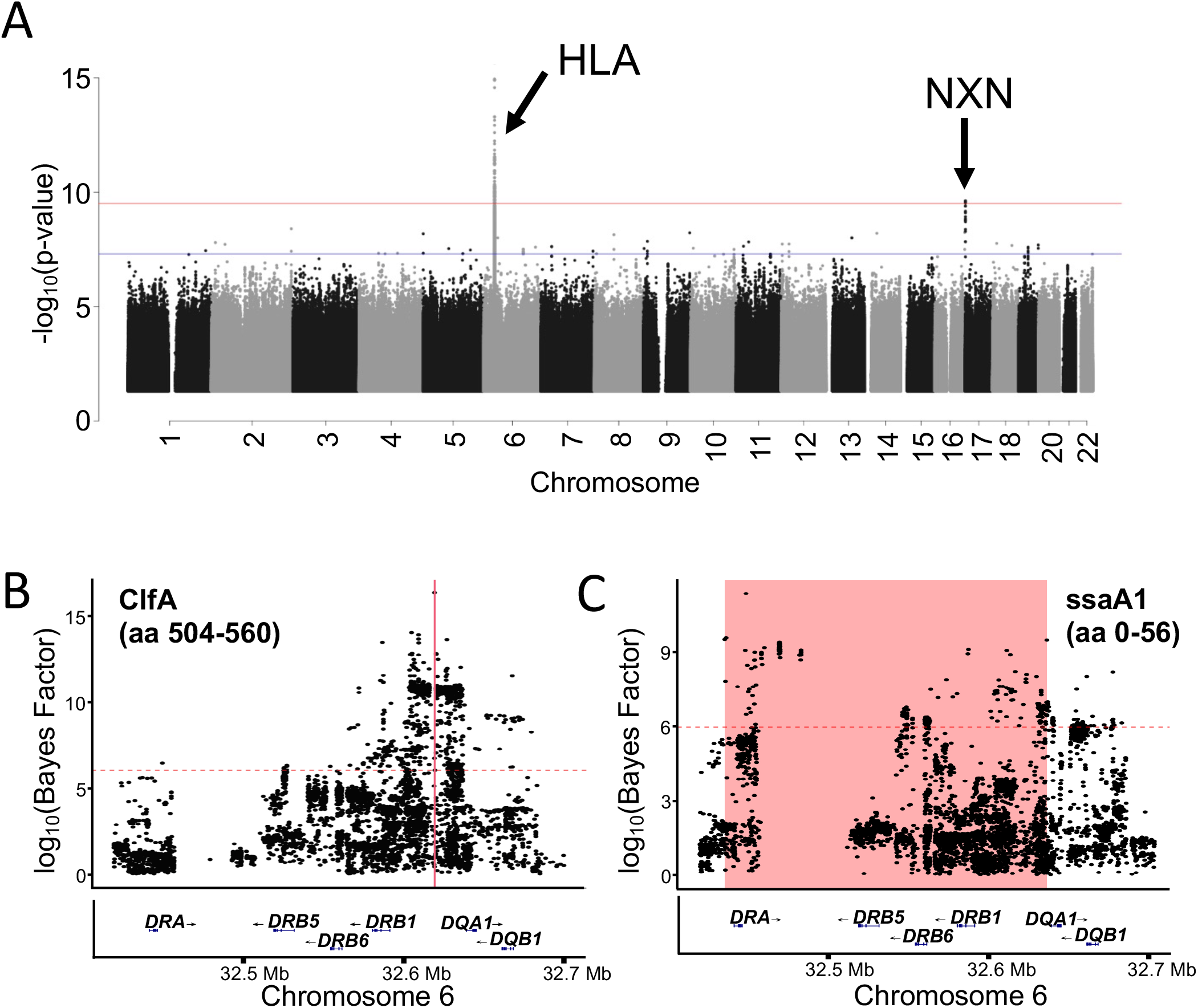
GWAS analysis of ToxScan reactivities in the VRC cohort. **A.** Three *Staphylococcus aureus* peptides (two overlapping peptides from clumping factor A protein and one from Staphylococcal secretory antigen) associated with the MHC-II locus on chromosome 6, and one peptide associated with nucleoredoxin gene *(NXN)* on chromosome 17. All variants with a log10 transformed Bayes factor of ≥ 6 (dashed red lines) were considered significant. **B.** Zoom plot for association with reactivity targeting two overlapping peptides from *S. aureus* clumping factor A (ClfA). Region defined by credible sets analysis shaded in pink. **C.** Similar to **B** but for *S. aureus* Staphylococcal secretory antigen A1 (ssaA1).

### Host phenotypic associations with ToxScan reactivities

We next sought to compare ToxScan reactivities between phenotypically grouped individuals, both at the peptide and protein level, using the PhIP-Seq casecontrol (phipcc) comparison software.^29^ As previously documented by others, we detected relatively increased reactivity to *Haemophilus influenzae* among males^30^ (**Fig. S4A-B**) and to *Chlamydia trachomatis* among females (**Fig. S4C-D**).^31^

In comparing ToxScan profiles from the healthy cohort to a cohort of patients with Crohn’s disease (**Table S4**), an inflammatory bowel disease, we detected relatively increased reactivity to several homologous flagellin peptides from a diverse set of bacteria. Flagellin, aside from mediating motility, contributes to bacterial invasion into and adhesion to host cells. Many flagellins also activate Toll-like-receptor 5 (TLR5), which is expressed on human intestinal epithelium and immune sentinel cells.^32^ Fourteen flagellin peptides were reactive in a significantly higher proportion of Crohn’s patients (**Fig. 6A**). Twelve of these fourteen peptides shared epitopes (**Fig. S5**).^33^ *Helicobacter mustelae* flagellin reactivity was detected in 60% of Crohn’s patients and only 4.2% of healthy individuals. Levels of reactivity to *H. mustelae* flagellin differed significantly between seropositive individuals in the two cohorts (Mann-Whitney U Test, p=0.003; **Fig. 6B**). We noted that among the seropositive Crohn’s patients, levels of reactivity to these peptides were not significantly associated with sex, age, Harvey-Bradshaw index of Crohn’s disease severity, or Montreal classification of disease location (F-test, p=0.78, **Fig. S6A-C**).

**Figure 6.**
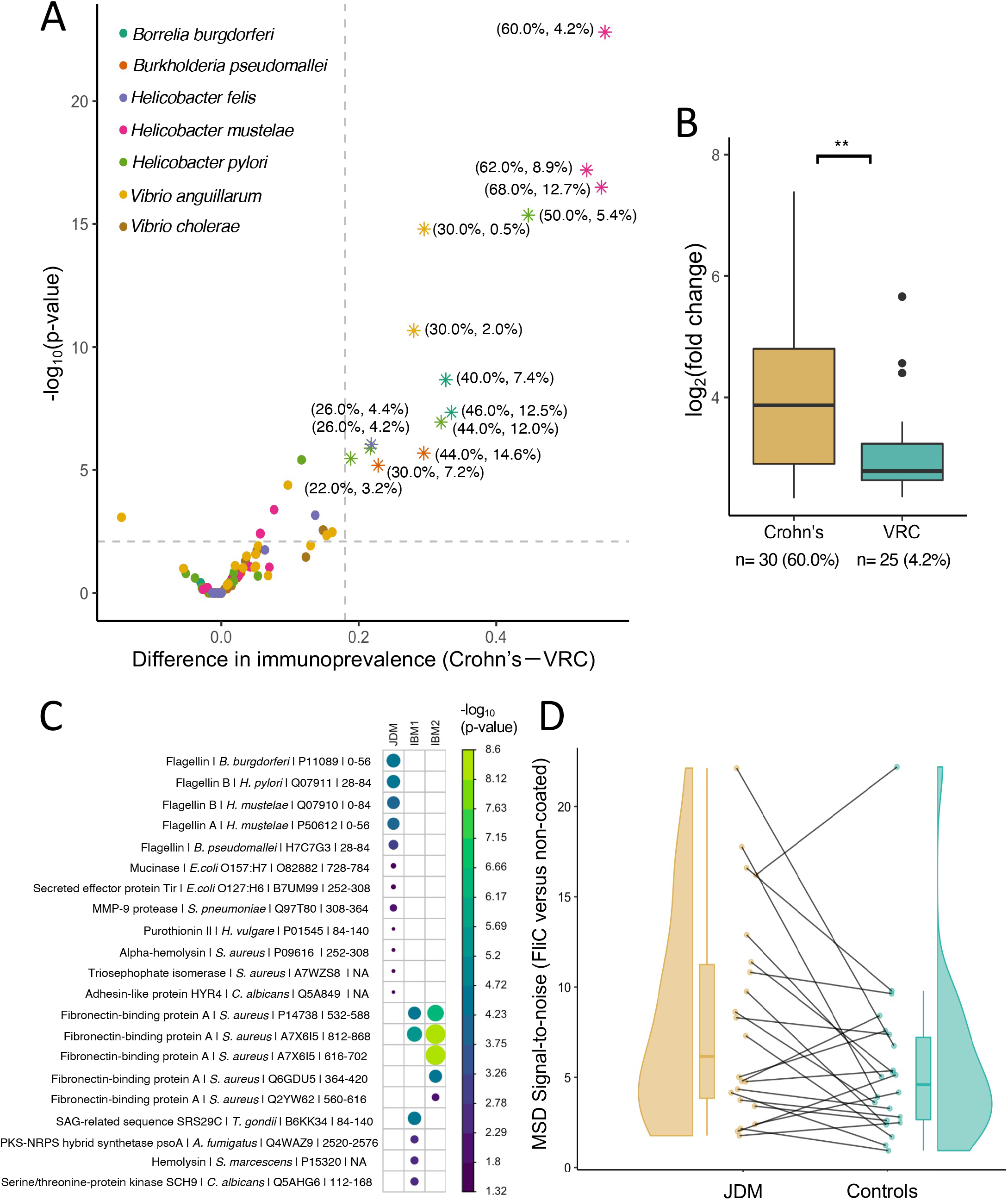
Association of ToxScan reactivities with autoinflammatory diseases. **A.** Volcano plot depicting differences in prevalence of antibody reactivities among Crohn’s disease patients versus the VRC cohort. Each point denotes a flagellin peptide. Significance of association was corrected for multiple testing using the Benjamini-Hochberg (BH) procedure. The 14 starred peptides are significantly associated with Crohn’s; these peptides share common epitopes (**Figure S5A**). The data labels indicate the percentage of the cohorts which are reactive (Crohn’s%, VRC%). **B.** Comparison of strengths of reactivity between the Crohn’s and VRC cohorts for the most differentially reactive peptide (*Helicobacter mustelae*, Mann-Whitney U Test, p=0.003). **C.** Correlation of ToxScan reactivities in 3 different myositis cohorts versus the VRC cohort (Fisher’s exact test with BH corrected p-values). **D.** flagellin FliC MSD ratio is shown for 22 JDM cases and their age-matched controls (matched pairs are connected by lines) using FliC antigen. Lines connect matched cases and controls.

### ToxScan profiles in idiopathic inflammatory myopathies

The etiologies of idiopathic inflammatory myopathies (IIMs) are believed to involve environmental factors.^34^ For example, low IIM disease concordances among monozygotic twins implicate strong contributions from the environment.^35,36^ We used ToxScan to profile serum samples obtained from the following cohorts: juvenile dermatomyositis (JDM, n=23, **Table S6**), seronegative inclusion body myositis (IBM) patients (IBM1, n=148, **Table S7**) and IBM patients with additional autoimmune or immunocompromised status (IBM2, n=57, **Table S7**). **Fig. 6C** describes the significance of myositis-ToxScan associations at the peptide level (compared to the VRC cohort). Interestingly, the two IBM cohorts shared significant associations with two homologous peptides from the *S. aureus* fibronectin-binding protein A. JDM, however, exhibits a different set of associations, the most significant being the same Crohn’s associated flagellin peptides.

Since JDM has not previously been associated with flagellin antibody reactivity, and because flagellin ELISA assays have been validated in the setting of Crohn’s disease, we sought to confirm the JDM reactivity using a Meso Scale Diagnostics (MSD) immunoassay. To this end, we assessed JDM serum antibody reactivity to the commonly studied *E. coli* flagellin FliC and compared it to that of a cohort comprised of same-gender siblings within five years of age (n=17) and unrelated age-matched controls (n=6). FliC reactivity was increased among JDM cases as compared to their age-matched controls (paired t-test, p = 0.03; **Fig. 6D**). Notably, of the 4 age-matched control samples with flagellin reactivity, all were siblings of flagellin seropositive JDM patients (1 was a monozygotic twin and 3 were non-twin siblings). Taken together, these findings suggest that immune responses to common microbial protein toxins/virulence factors may contribute to IIM disease pathogenesis.

We further investigated additional characteristics of JDM patients with and without flagellin reactivity. JDM cases with flagellin reactivity were diagnosed at an earlier age (≤10 years old, p = 0.037). Interestingly, 9 of the 11 (82%) JDM samples with flagellin reactivity were also transcriptional intermediary factor 1 gamma (TIF1-γ) autoantibody positive, versus only 5 of the 12 (42%) without flagellin reactivity. Flagellin reactivity may therefore serve as a novel biomarker that defines a distinct JDM endotype.

## Discussion

We have designed and constructed a novel programmable phage display library, ToxScan, to represent the 14,430 non-viral environmental and microbial toxins and virulence factors associated with 1,312 genera, which were present in the UniProt database in November 2016. The ToxScan library is a renewable resource that can be used for unbiased investigations associating health or disease states with immune responses to environmental protein toxins and virulence factors. Here we employed PhIP-Seq with the ToxScan library to profile serum IgG antibody reactivities in crosssectional and longitudinal cohorts of healthy individuals and in patients with Crohn’s disease or idiopathic inflammatory myopathy.

In a cohort of 598 healthy adult volunteers (the VRC cohort), we detected 848 frequently reactive peptides (>5% seroprevalence). This set of immunoprevalent peptides includes at least 364 non-overlapping epitopes; 549 of these 848 peptides share no significant homology with sequences previously reported in the Immune Epitope Database, IEDB. We performed a focused analysis of antibody reactivities to immunoprevalent peptides derived from cell surface and secreted proteins of *S. pneumoniae, S. pyogenes, S. aureus, S. epidermidis, E. coli*, and *H. influenzae*, observing pronounced variation across subgroups of individuals, likely reflecting differential exposure or colonization profiles, and protection or susceptibility to future infections.

We observed prevalent recognition of the homologous pneumococcal histidine triad proteins D and E (PhtD and PhtE). PhtD is produced by all pneumococcal strains and has been explored as a candidate vaccine antigen.^37^ Pathogenic *S. pneumoniae* rely on Pht proteins for lung epithelial cell adhesion, which can be inhibited by antibodies that bind PhtD.^38^ In mouse models, Pht immunization elicits protection against pneumonia, sepsis,^39^ and nasopharyngeal colonization.^40^ C3 complement deposition increases upon deletion of PhtA, PhtB, PhtD, and PhtE, suggesting that these proteins may also play a role in evading complement-dependent defenses.^41^ Additional studies will define the association between the prevalent Pht antibodies we detected using ToxScan and their protective activities against tissue-invasive *S. pneumoniae*.

We detected subgroups of individuals with increased IgG responses to *S. pyogenes* C5a peptidase (SCPA) and M protein serotype 5 (M5). C5a peptidase participates in host immune evasion by inactivating both C3a and C5a, two complement proteins that act as chemoattractants to recruit neutrophils to sites of infection.^42^ M5, M1, and M6 (but not M4, M22, or M60) are able to bind human fibrinogen, forming a complex that can contribute to host evasion by inhibiting human C3 deposition on *S. pyogenes*.^43^ It will be interesting to determine whether the antibodies identified in this study can neutralize these mediators of immune evasion. Antibodies to Streptococcal M proteins can cross-react with host proteins including myosin, and are thought to participate in post-streptococcal autoimmune sequelae, such as rheumatic fever associated myocarditis. The relationship between those antibodies and the specificities detected with ToxScan remain to be determined.^44^

We detected 99.5% seropositivity for at least one of the *S. aureus* peptidoglycan hydrolases, the N-acetylmuramoyl-L-alanine amidase Sle1 and the bifunctional autolysin Atl. *S. epidermidis* Sle1 reactivity was observed in the same individuals, likely due to a high degree of sequence similarity to *S. aureus* Sle1 (75.5% identity).^45^ Loss of peptidoglycan hydrolase activity interferes with dissemination of *S. aureus* and thus decreased virulence in a murine model.^46^ Host antibody responses that interfere with peptidoglycan hydrolase activity may therefore diminish the efficiency of staphylococcal tissue invasion.^47^ The small subset of individuals who appear to lack these antibodies may therefore be at greater risk of more serious staphylococcal infections.

We detected near universal reactivity to the *E. coli* translocated intimin receptor (Tir), most likely driven by the common exposure in childhood to enteropathogenic (EPEC) strains.^48^ EPEC is a common cause of diarrhea in children while enterohemorrhagic (EHEC) strains are associated with foodborne outbreaks in the United States. In pediatric patients with *E. coli* O157:H7 (EHEC) infection, antibodies to Tir were more prominent versus other virulence factors.^49^ Immunization of calves with intimin, Tir, and secreted protein A resulted in decreased shedding of *E. coli* O157:H7 after challenge, suggesting a potentially protective role for these antibodies.^50^ These component vaccines elicit antibody cross-recognition of different *E. coli* serotypes, likely due to targeting of functional, conserved epitopes.^51^

We detected prevalent, yet less immunodominant reactivity to the *H. influenzae* immunoglobulin A1 protease (IgAP) autotransporter. IgAP is of a particular research interest with regard to nontypeable *H. influenzae*, which is thought to contribute to chronic obstructive pulmonary disease (COPD) if present in the lower airways.^52^ IgAP is an endopeptidase that proteolytically inactivates IgA to evade host immune response. Fragments of cleaved IgA have been detected in the sputum of patients with COPD^53^; the association of the IgAP antibodies described here and these IgA fragments, along with the risk for developing COPD, is therefore worthy of further investigation.

We next used a longitudinal cohort (BLSA) to assess the stability of ToxScan reactivity profiles. We observed that the intra-individual variability of the number of ToxScan reactive peptides was far lower than the inter-individual variability, suggesting that individuals establish a relatively stable environmental antibody diversity set point in adulthood. We found this set point to slowly decline with age by ~0.89 reactivities per year, which could be attributed to a multiplicity of factors. It could reflect age-related immunosenescence, which is associated with loss of B cell production, immunoglobulin diversity, and responsiveness to vaccination.^54^ An alternative explanation is that the diversity of antibodies to environmental antigens continues to expand over time, thereby diluting the concentration of the individual antibody specificities detectable at earlier timepoints.

Our longitudinal study also revealed that the antibody specificities acquired by adulthood tend to persist over time. Non-heritable factors that influence epitope selection are likely diverse, and include health status at the time of exposure, prior immune responses, the size of the inoculum, and whether or not a tissue-damaging infection is established. The contribution of heritable factors to antibody epitope selection has been investigated in the context of viral infection,^55,56^ but not yet in the context of anti-toxin or anti-virulence factor antibody epitope selection. Multiple peptide reactivities mapped to the MHC-II region of the HLA locus on chromosome 6, while one peptide mapped to a non-HLA locus on chromosome 17. The majority of antibody reactivities were not significantly associated with any genomic locus, but this could be due to the relatively low power of our study. Even robust associations often require thousands of individuals for genome-wide trait mapping. We expect that future, larger studies will identify many more MHC-II associated ToxScan reactivities, as well as pathogen response-related loci outside of HLA, potentially elucidating novel mechanisms of host-microbe interactions.

We compared ToxScan profiles from the cross-sectional VRC cohort to profiles from a Crohn’s cohort. This analysis detected the well-known Crohn’s-associated bacterial flagellin antibody reactivity. Of the fourteen flagellin peptides more reactive in Crohn’s patients, twelve shared the conserved epitope (**Fig. S5**, aa: KLSSGLRINKAADDASGMAIA). This sequence was also shared (100% sequence identity) by the commonly studied flagellin protein, *E. coli* serotype 044:H18 FliC. ^57,58^ The most differentially reactive peptide was from a *H. mustelae* flagellin protein, which shares 82.1% sequence similarity with the most differentially reactive *H. pylori* flagellin peptide. Intriguingly, prior studies have found a lower seroprevalence of *H. pylori* in Crohn’s patients compared to healthy controls.^59,60^

Interestingly, we also found that bacterial flagellin reactivity was associated with juvenile dermatomyositis (JDM) patients compared to matched controls. This finding complements a previously described anti-myosin autoantibody, thought to be associated with response to *S. pyogenes* M5 protein.^61^ Strikingly, all 11 flagellin positive JDM patients were younger than age 10, with the majority of them (9/11) also being TIF1-γ autoantibody positive. TIF1-γ^62,63^ is found in 18-30% of JDM patients and is associated with severe cutaneous disease and a chronic disease course. These observations suggest a specific association of flagellin seropositivity with a young, TIF1-γ positive subset of JDM patients.

Since flagellin immunoreactivity is considered a biomarker for Crohn’s disease, we performed a medical record review for gastrointestinal symptoms in the flagellin seropositive JDM patients, which failed to detect an increase in gastrointestinal symptoms. Of the four JDM age-matched controls who were flagellin positive, all were siblings (three non-twin siblings and one monozygotic twin) of flagellin positive JDM patients. Since the majority of the age-matched controls were non-twin siblings, and since a much higher fraction of the sibling controls were flagellin positive compared to the VRC cohort, it is likely that heritable or common environmental factors strongly influence the likelihood of developing flagellin antibodies. If flagellin antibodies are causally related to the development of JDM, however, they are related with incomplete penetrance. These results argue for additional, more comprehensive analyses of the microbiota and corresponding immune responses in JDM.

The technical limitations of programmable phage display-based antibody profiling also apply to the ToxScan library. Here we have utilized 200-mer oligonucleotide library synthesis to encode 56-aa peptides. This length does accommodate some degree of secondary structure, but highly conformational and discontinuous epitopes will be absent from the library. The phage-displayed peptides will also lack post-translational modifications, which frequently contribute to epitopes as haptens. Antigenic representation in the library is further limited by the accuracy and completeness of existing protein annotations, and the removal of highly similar sequences during library design. Furthermore, virulence factors and toxins may be associated with multiple organisms and the specificity of the linkage to these organisms may be incompletely defined. This latter consideration should inform the interpretation of results like the association of *H. mustelae* flagellin antibodies with Crohn’s disease, since *H. mustelae* is primarily found in ferrets. Finally, the typical size of synthetic oligonucleotide libraries (~10^5^) constrains our ability to represent all components of the microbiota. Future library design paradigms will expand the antigenic spaces that can be represented.

In this study, we have adapted the standard PhIP-Seq assay protocol to ToxScan profiling, which involves protein A/G coated magnetic bead immunocapture. This approach primarily captures IgG molecules, with potentially minor contributions from IgA and IgM binding to protein A. However, in many settings, particularly those involving mucosal immunity, IgA is the dominant antibody isotype. For such projects, IgA-specific ToxScan analyses may therefore be more appropriate.

While this manuscript was in preparation, Vogl et. al. reported a PhIP-Seq analysis of 997 healthy individuals using a 244,000 peptide library that includes surface and secreted proteins from commensal, pathogenic and probiotic bacterial species, as well as the Virulence Factor Database and additional controls from IEDB.^22^ Similar to our findings, the Vogl study detected antibodies to many public and personal epitopes from both pathogenic and commensal bacteria. These reactivities were stable over time and correlated weakly with metagenomic sequencing of the same individuals. Because PhIP-Seq libraries are modular in nature, The Vogl and ToxScan libraries could be combined and screened simultaneously. Further combining them with the VirScan library, the AllerScan library, and even the human proteome library, would provide an unprecedented wealth of information useful for dissecting the complex interplay between environmental exposures, commensal colonization, infections, and host disease pathogenesis.

## Materials and Methods

### Design and cloning of ToxScan peptide library

Design and cloning of the ToxScan peptide library followed previously published PhIP-Seq protocols.^16^ In November 2016, we downloaded all protein sequences from the UniProt database annotated with keywords, “toxin” (12,679 proteins representing clusters at >90% sequence identity) and “virulence factor” (all 2,133 unique proteins). The pepsyn^16^ library design software converted the 14,430 unique protein sequences into 95,601 overlapping 56 amino acid peptide tiles with 28 amino acid overlaps, which were reverse-translated into their corresponding DNA sequences. Silent mutations were made to remove EcoRI and XhoI restriction sites, which were utilized for cloning. PCR primer binding sequences AGGAATTCCGCTGCGT and CAGGGAAGAGCTCGAA were added to the 5’ end and the 3’ end, respectively, bringing the total oligonucleotide length to 200 bases. These encoding oligonucleotides were synthesized (Twist Bioscience) and cloned into the mid-copy T7-FNS2 phage display vector as described previously,^17^ and packaged using the T7 Select Packaging Kit (EMD Millipore). Library quality assessment was performed as described in Results.

### Study participants

#### VRC cohort

Plasma was collected from 598 healthy volunteers who participated in the National Institutes of Health (NIH) Vaccine Research Center’s (VRC)/National Institutes of Allergy and Infectious Diseases (NIAID)/NIH protocol “VRC 0θO: Screening Subjects for HIV Vaccine Research Studies” (NCT00031304) in compliance with NIAID IRB approved procedures, including informed consent. The demographic composition of this cohort is provided in **Table S1**. CMV serostatus of the participants was determined using VirScan data and the AVARDA algorithm.^10^

The following quality-control criteria were employed for sample inclusion: reactive to at least 0.1% of the ToxScan library (>100 peptides total) and reactive to at least 1% of the immunoprevalent peptides (>10 immunoprevalent peptides).

#### BLSA cohort

Sera were obtained from 47 healthy volunteers at two timepoints roughly 5 years apart as part of the Baltimore Longitudinal Study of Aging (BLSA) at the National Institute of Aging (NIA). Clinical metadata regarding age and timepoints are displayed in **Table S3**.

#### Crohn’s cohort

Sera were obtained from 50 Crohn’s disease patients as part of the IRB-approved initiative, Stratifying Medication And Refractory to Therapy IBD (SMART-IBD) at the Jill Roberts Center of IBD at Weill Cornell in New York. 25 patients were being treated for L1: ileal Crohn’s disease while the remaining 25 were being treated for L3: ileocolonic Crohn’s disease according to the Montreal classification for inflammatory bowel disease. Additional demographic metadata are provided in **Table S4**.

#### JDM cohort

Sera from 23 juvenile dermatomyositis (JDM) patients meeting EULAR-ACR Myositis Classification Criteria^64^ and 23 age-matched controls were collected as part of ongoing natural history studies at the NIH (NCT00017914 and NCT00055055) (**Table S5**).

#### IBM cohorts

Sera from 205 inclusion body myositis (IBM) patients were obtained as part of an ongoing observational study at the Johns Hopkins Myositis Center. IBM patient samples were collected under protocol IRB00235256. All patients met ENMC 2011 diagnostic criteria and provided informed consent. These IBM patients’ sera were separated into two cohorts: those with other autoimmune and/or immune-mediated diseases (57 patients) and those who were seronegative for myositis associated antibodies (148 patients).

### Phage ImmunoPrecipitation Sequencing

The standard protein A/G PhIP-Seq protocol, which was employed here, has been described in detail.^16^ Briefly, ELISA was performed to measure total IgG in serum samples and input volume was adjusted to 2 μg of IgG input per immunoprecipitation (IP). The ToxScan and VirScan libraries were mixed with diluted serum at 10^5^-fold library coverage. The mixture was allowed to rotate overnight at 4°C, followed by a 4hour IP with protein A and protein G coated magnetic beads (Dynal beads, Invitrogen). A first PCR was performed with primers that flank the displayed peptide inserts. A subsequent PCR added adapters and indexes for Illumina sequencing. Fastq files were aligned to obtain read count values for each peptide in the library, followed by detection of antibody reactivity by comparison with a set of mock IPs run on the same plate.^65^

For a peptide to be considered reactive, the counts needed to exceed 15, the maximum likelihood fold change needed to be at least 5, and the p-value of differential abundance needed to be lower than 0.001 (fold-changes and p-values were calculated using the EdgeR software). All heatmaps were constructed using the pheatmap package and the tidyverse package suite. Heatmaps were clustered according to hierarchical analysis as computationally determined by Ward’s method (Ward.D2) and the Euclidean distance measure. All network graph analyses utilized the igraph package in R.^66^

We used a one-way ANCOVA to assess whether there was an association between the diversity of ToxScan reactivity and sex, age, race, or CMV status (**Fig. 4A**). Volcano plots were corrected for multiple comparisons using the Benjamini-Hochberg procedure.

### Meso Scale Diagnostics (MSD) testing for flagellin antibodies

MSD testing of the flagellin antibodies was performed according to the manufacturer’s guidelines.^67^ Recombinant FliC was prepared as a 6X-His-tagged protein in *E. coli* BL21 and purified by metal affinity chromatography as described.^68^ FliC was diluted to a concentration of 6.8 μg/mL in PBS. Briefly, MSD 96-well MESO QuickPlex SQ 120MM Standard plates (cat #: L55XA) were incubated with 25 μL of diluted FliC antigen per well, overnight at 4°C. Another plate, which served as the “uncoated” negative control, was incubated overnight with only PBS at 4°C. Subsequent steps were performed the same way for both the FliC coated plate and the uncoated plate. Plates were then blocked overnight at 4°C in 150 μL of blocking buffer per well (1 % w/v MSD blocker A, cat #: R93BA-4) in PBS. Plates were washed 5 times in 0.05% PBS-T. Sera were diluted 1: 10,000 in blocking buffer and 25 μL of each diluted serum sample were added to the plates. The plates were incubated at room temperature for 1.5 hrs with shaking then washed 5 times with 0.05% PBS-T. 25 μL of 1:500 detection antibody (MSD SULFO-TAG goat anti-human antibody, cat #: R32AJ-5) diluted in blocking buffer was added to each well and the plates were incubated at room temperature for 1.5 hrs with shaking. Plates were washed again 5 times with 0.05% PBS-T. 150 μL of MSD GOLD read buffer (cat #: R92TG-2) was added to each well. Plates were read with a MSD QuickPlex SQ 12 Instrument to determine an intensity value for each well on each plate. FliC dependent signals (“MSD ratios”) were determined for each sample by dividing the MSD intensity value from the FliC coated well by the corresponding intensity value from the uncoated well.

### GWAS of ToxScan epitope selection

After binarization of the ToxScan peptide reactivity data, we considered only the 166 peptides with seroprevalence between 16% and 80% in the VRC cohort (**Fig. S2**). Genotyping of the VRC cohort and imputation of genetic variants are described in detail elsewhere.^55^ We interrogated 7,637,921 variants (imputed from 2,783,635 genetic variants with a minor allele frequency ≥ 5%, measured using the Illumina Human Omni 5 BeadChip array, GRCh37) for an association with each of the 166 ToxScan peptides using the penalized quasi-likelihood (PQL) approximation to the GLMM (Breslow and Clayton) implemented in the R package Genesis.^69,70^ Principal component analysis was utilized to determine most likely ancestry of the individuals rather than self-reported demographic characteristics. The African ancestry GWAS included the genetic relationship matrix (GRM PC-Relate) as a fixed effect and 10 principal components as random effects. We first performed single-variant association analysis using the VRC donors of European ancestry (VRC/EUR n=478); replication was then performed using the VRC/AFR cohort (n=147). The significance threshold for discovery was estimated with the Sidak-Nyholt method,^71,72^ accounting for the number of independent traits (n=161.7) resulting in a genome-wide significance for ToxScan peptides of p-value ≤ 3.09×10^-10^.

Credible sets analysis was performed as described previously.^55^

### Correlation plots

The 1,000 most seroprevalent peptides (up to 80%) from the VirScan and ToxScan libraries were used to identify a maximal vertex set of independent peptide reactivities based on sequence similarity (iGraph R package). A group of 195 peptides were selected from the VirScan library and 173 were analogously selected from the ToxScan library. We calculated a Pearson’s R-value correlating the peptide fold changes among every sample pair and plotted them using the “corrplot” R package.^73^ Samples were quality-controlled utilizing the following criteria. If a pair of pre/post samples was not correlated in their ToxScan or VirScan reactivities (had a Pearson’s correlation coefficient below 0.7) and the VirScan profile was highly correlated with another sample besides its pair, the samples were assumed to be mislabeled and removed.

Myositis association p-values were determined by PhIP-Seq case-control (phipcc) software.^29^ Proteins in the correlation plot are those that were also significant when myositis samples were compared to control samples from the same PhIP-Seq plates, reducing the likelihood that significant hits were due to experimental batch effect.

### Network Graph and BLAST Analysis

To evaluate the degree of interconnectedness within the 848 immunoprevalent peptides, we constructed an undirected network graph using R igraph software package. To define peptide-peptide connectivity, we performed a blastp alignment, reporting all peptide-peptide alignments with a bitscore > 30 and an E value < 1. Each node in the network represents a peptide and each edge represents a blastp peptide-peptide alignment. We additionally evaluated overlap between the 848 immunoprevalent peptides and the 245,615 linear, B-cell derived, human-host epitopes present in the Immune Epitope Database (IEDB), by performing a blastp alignment with the same thresholds as as above. Blastp parameters were: “-evalue 1-max_hsps 1 -seg no -comp_based_stats none -soft_masking false -word_size 7 -max_target_seqs 100000”. We additionally used the igraph *independence.number* function to determine the largest set of non-overlapping peptides within the 848 immunoprevalent peptides. For the organism-specific homology analyses, the immunoprevalent peptides were compared with 245,615 IEDB epitopes, subsetted to specific organisms as indicated (*Streptococcus pneumoniae*, *Streptococcus pyogenes*, *Staphylococcus aureus, Staphylococcus epidermidis, Escherichia coli*, and *Haemophilus influenzae*, respectively).

## Acknowledgements

We thank Drs. Terrance O’Hanlon and Frederick Miller for their support related to the juvenile dermatomyositis samples. We thank Drs. Luigi Ferrucci, Madhav Thambisetty, and Vijay R. Varma for their support related to the BLSA samples. We thank Drs. Peter Burbelo and Sandra Wolin for their critical reading of the manuscript. We are grateful to Stephen J. Elledge for generously providing the VirScan library used in this study.

## Author Contributions

J.W.A., D.R.M., A.C., and H.B.L. conceptualized the study. U.L. and H.B.L. designed the ToxScan library. J.W.A. and H.B.L. wrote the original draft of the manuscript. J.W.A., D.R.M., A.C., T.V., and S.J. performed formal analysis. D.R.M. and B.M.S. performed PhIP-Seq testing and library screening, and developed software for PhIP-Seq data analysis. C.V., T.V., and P.D. performed the GWAS analysis. S.J. and D.R.M. performed the MSD assays and analysis using reagents provided by M.Q.W. M.R. and T.L. provided VRC samples, R.S. and C.A.S. provided BLSA samples, L.G.R. and P.N.F. provided JDM and age-matched healthy control samples, T.E.L. and A.L.M. provided IBM samples, and R.S.L. and G.F. provided Crohn’s samples. T.S.S., C.L.S. and P.J.S. reviewed the manuscript and provided scientific input. H.B.L supervised the project.

## Supplementary Figure Captions

**Figure S1.**
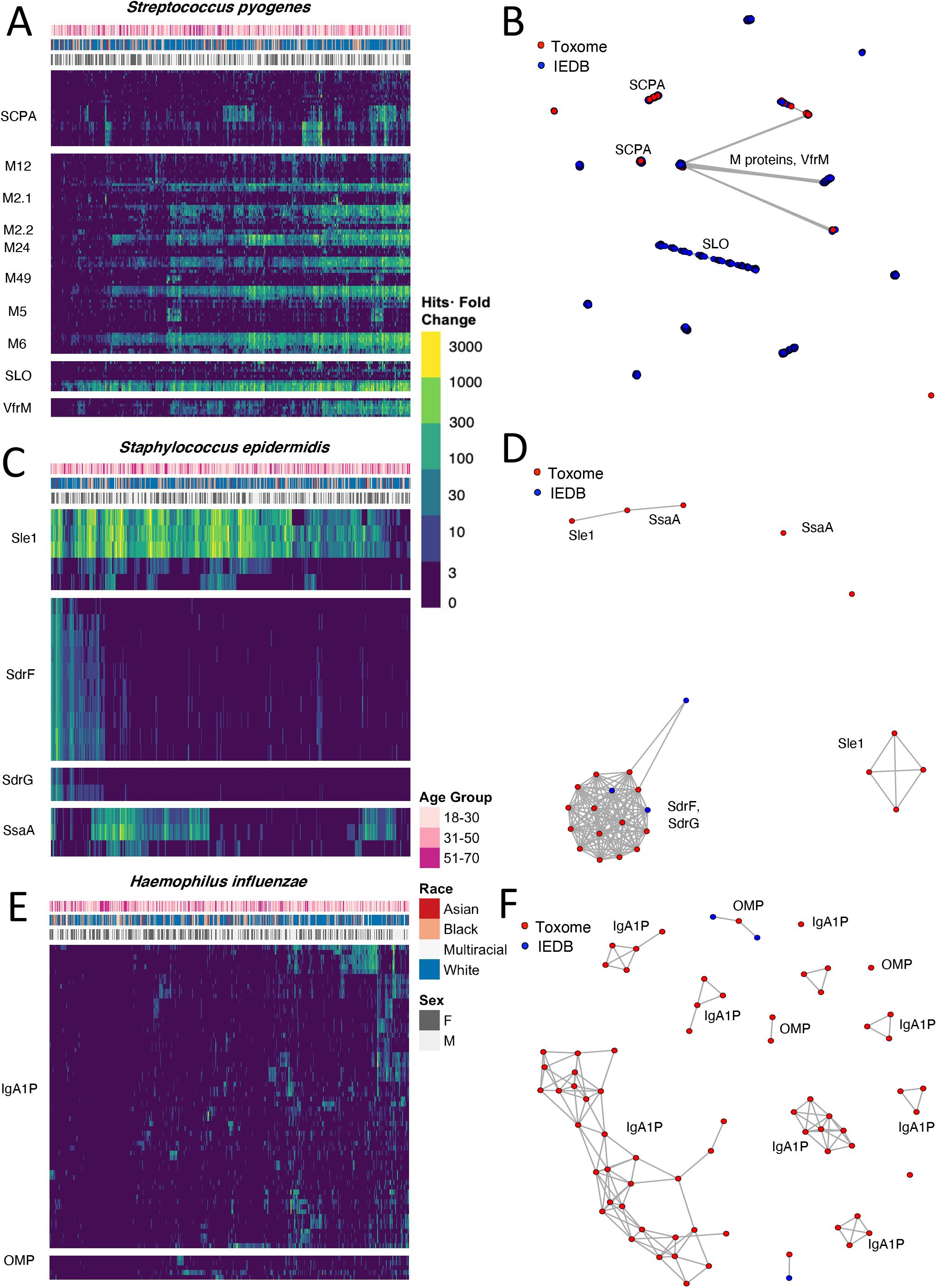
Immunoreactivity targeting proteins from three additional commonly encountered bacteria. **A.** Reactivities to *S. pyogenes:* C5a peptidase (SCPA), M protein serotypes 12, 2.1,2.2, 24, 49, 5, and 6, streptolysin O (SLO), and virulence factor-related M protein (VfrM). ***B***. Shared epitope network graph depicting *S. pyogenes* immunoprevalent ToxScan peptides and linear IEDB antibody epitopes. **C.** Reactivities to *S. epidermidis:* N-acetylmuramoyl-L-alanine amidase Sle1, serine-aspartate repeat containing proteins F and G (SdrF, SdrG), and Staphylococcal secretory antigen SsaA. **D.** Shared epitope network graph depicting *S. epidermidis* immunoprevalent ToxScan peptides and linear IEDB antibody epitopes from *S. aureus*. **E.** Reactivities to *H. influenzae:* immunoglobulin A1 protease (IgA1P) and outer membrane protein (OMP). **F.** Shared epitope network graph depicting *H. influenzae* immunoprevalent ToxScan peptides and linear IEDB antibody epitopes.

**Figure S2.**
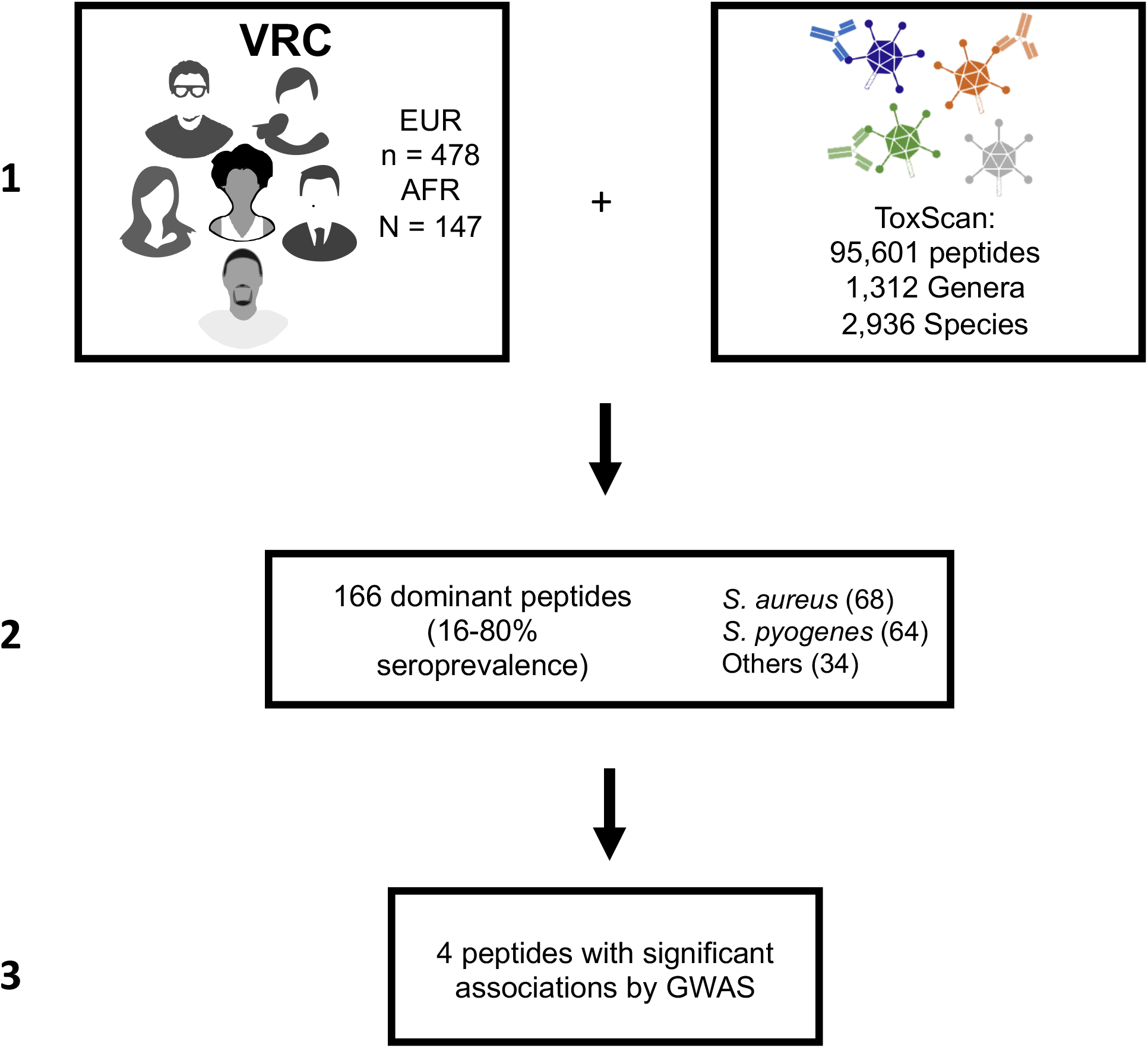
Flow chart of ToxScan peptide selection for GWAS. Peptide reactivity scores were binarized (responders and non-responders) and filtered based on responder proportion (≥16% and ≤80%) in the VRC cohort. The filtered 166 immunoprevalent peptides were subjected to genome-wide association study (GWAS) in the VRC/EUR subgroup. 4 peptides exhibited significant genome-wide associations.

**Figure S3.**
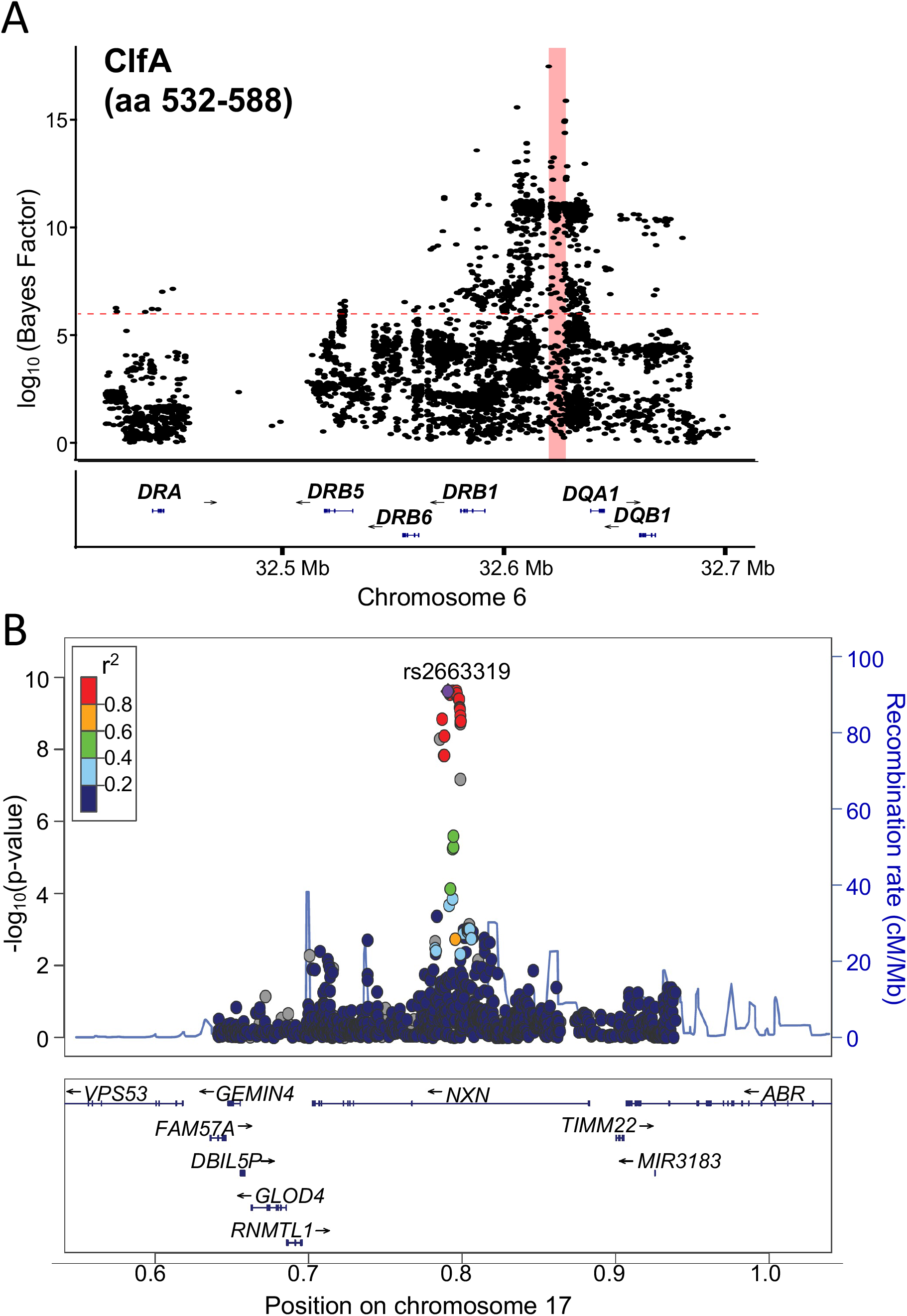
**A.** Locus zoom plot of CLFA (532-588), confirming findings from adjacent peptide CLFA (504-560). The region in pink was identified by credible sets analysis. **B.** Locus zoom plot (Chr 17) for *Streptococcus pyogenes* serotype M3 Streptolysin O peptide.

**Figure S4.**
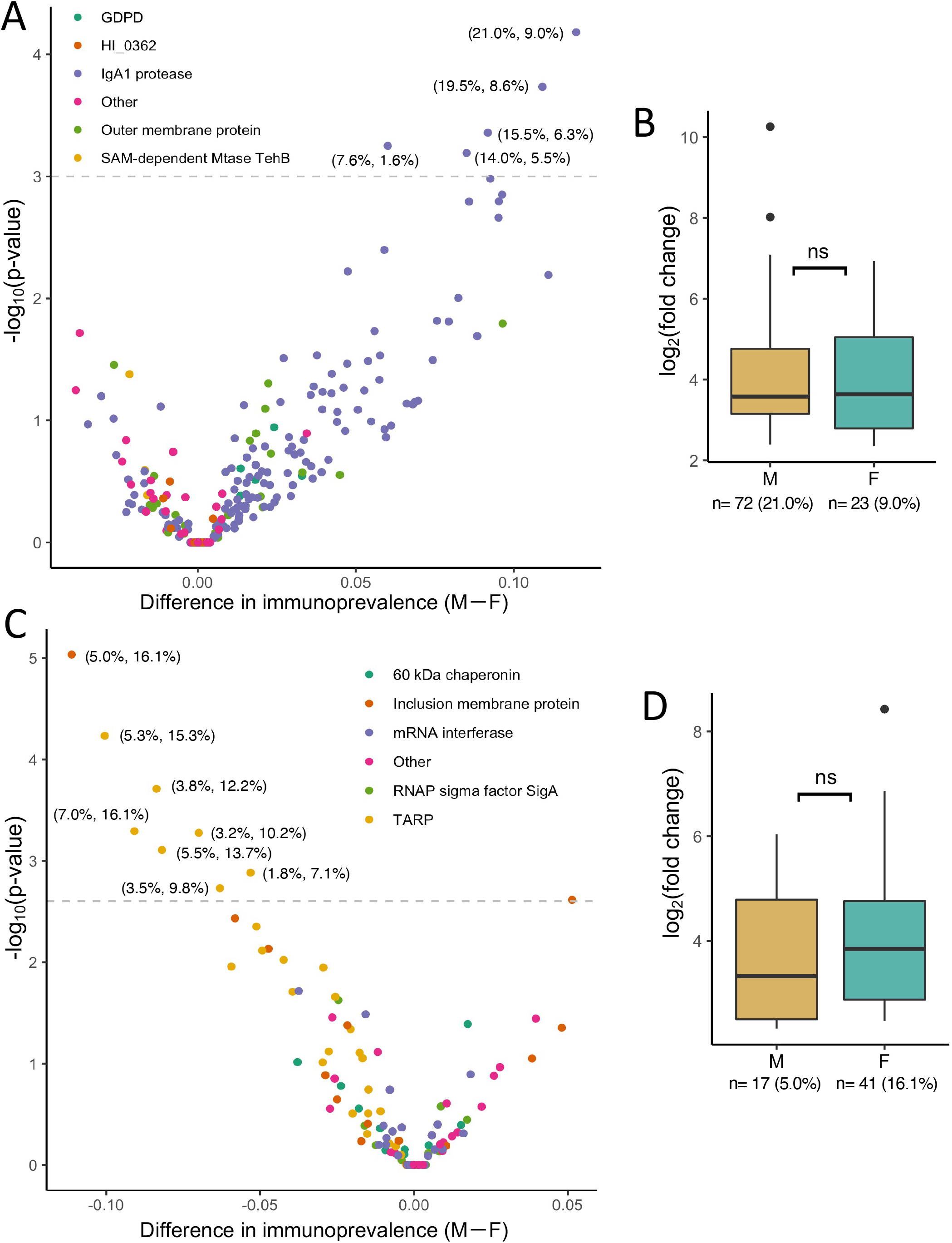
Differential reactivities observed between males and females in the VRC cohort. **A.** Each point represents a *H. influenzae* virulence factor or toxin peptide. Reactivities plotted: glycerophosphodiester phosphodiesterase (GDPDL), uncharacterized periplasmic iron-binding protein HI_0362, immunoglobulin A1 protease autotransporter (IgA1), outer membrane protein, and probable S-adenosyl-L-methionine-dependent methyltransferase TehB (SAM-dependent MTase TehB). Peptides above the significance threshold are more frequently reactive in males (from IgA1 protease). The data labels indicate immunoprevalence by sex (% female, % male) **B.** The strength of the most significantly male-specific *H. influenzae* reactivity (from IgA1 protease) among seropositive individuals by sex (no difference). **C.** Each point represents a *Chlamydia trachomatis* virulence factor or toxin peptide. Reactivities plotted: 60 kDa chaperonin, inclusion membrane protein, mRNA interferase, RNA polymerase (RNAP) sigma factor SigA, and translocated actin-recruiting phosphoprotein (TARP). Peptides above the significance threshold are more frequently reactive in females (from TARP and inclusion membrane protein). **D.** The strength of the most significant female-specific *Chlamydia trachomatis* reactivity (from inclusion membrane protein) among seropositive individuals by sex (no difference).

**Figure S5.**
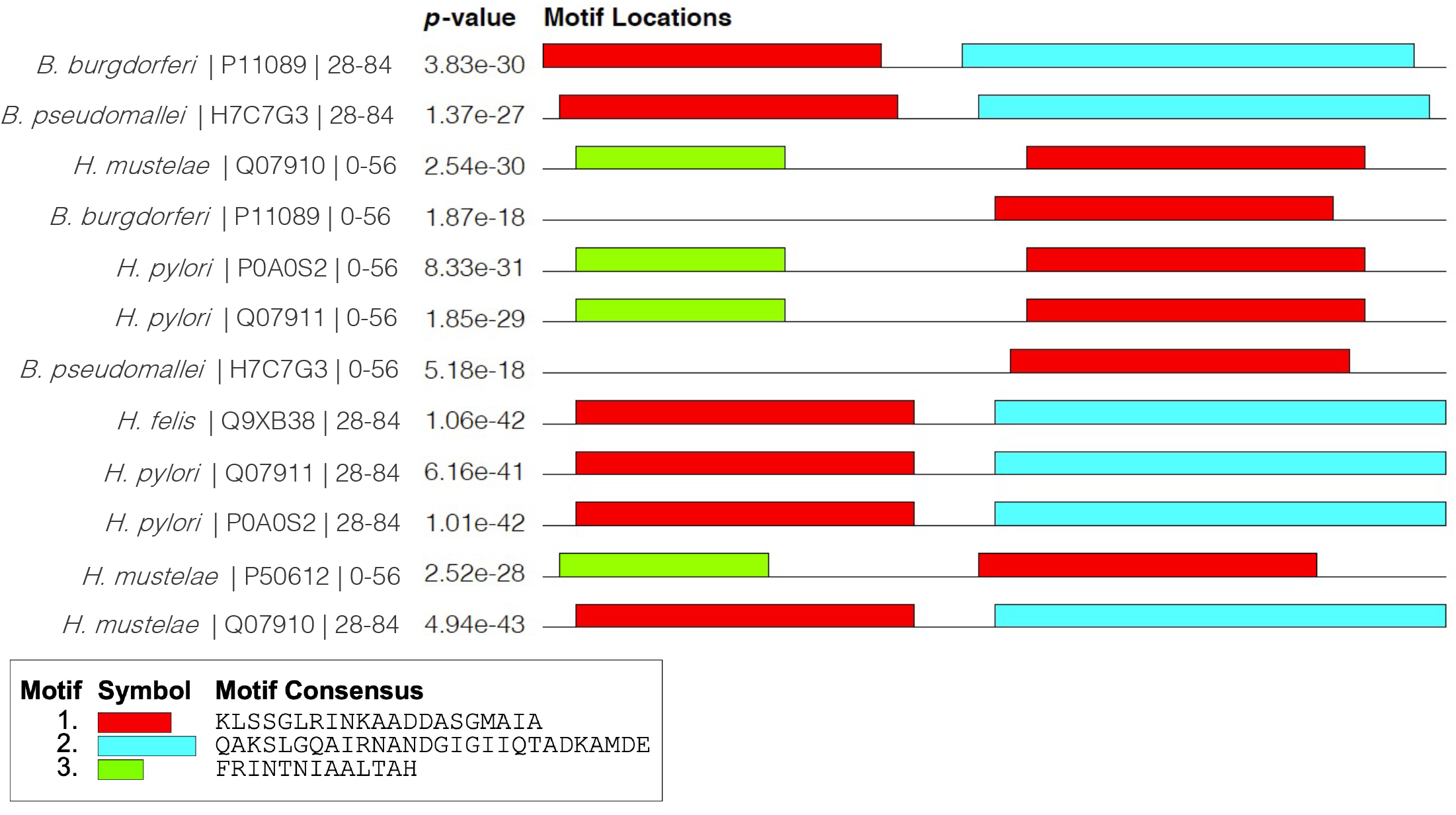
Shared motifs among Crohn’s reactive flagellin peptides. Twelve of the 14 flagellin peptides more reactive in Crohn’s patients versus the VRC cohort share common epitopes. Results from the MEME Suite analysis are shown.

**Figure S6.**
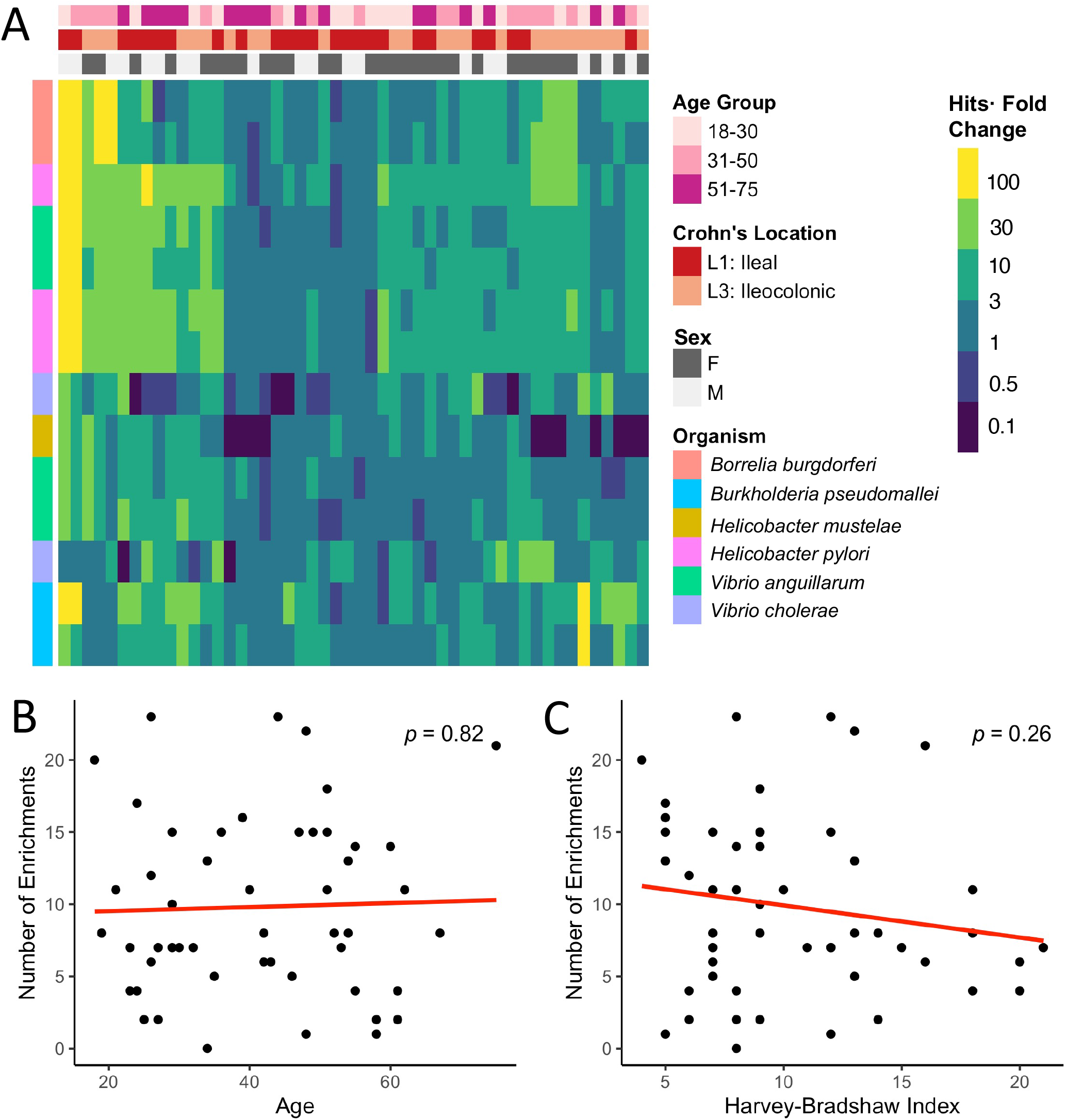
Lack of phenotypic association with flagellin reactivity among Crohn’s patients. **A.** Reactivity of 14 Crohn’s specific flagellin peptides (starred peptides in **Fig. 6A**). Each row is a peptide and each column is a sample. No correlation was detected between number of Crohn’s associated flagellin peptide reactivities and age (**B**) or Harvey-Bradshaw severity index (**C**).

## Supplementary Table Captions

**Table S1.** Demographic information for the VRC Cohort. Table lists sex, age at time of sampling, race, and CMV status.

| Study population (n=598) |  |
| --- | --- |
| SEX |  |
| Male | 343 (57.4%) |
| Female | 255 (42.6%) |
| AGE (in years) |  |
| Mean | 34.8 ± 12.9 |
| Median | 31 |
| Range | 18–70 |
| RACE |  |
| Asian | 23 (3.8%) |
| Black | 152 (25.4%) |
| Multiracial | 21 (3.5%) |
| White | 402 (67.2%) |
| CMV STATUS |  |
| Seropositive | 295 (49.3%) |
| Seronegative | 303 (50.7%) |

**Table S2.**
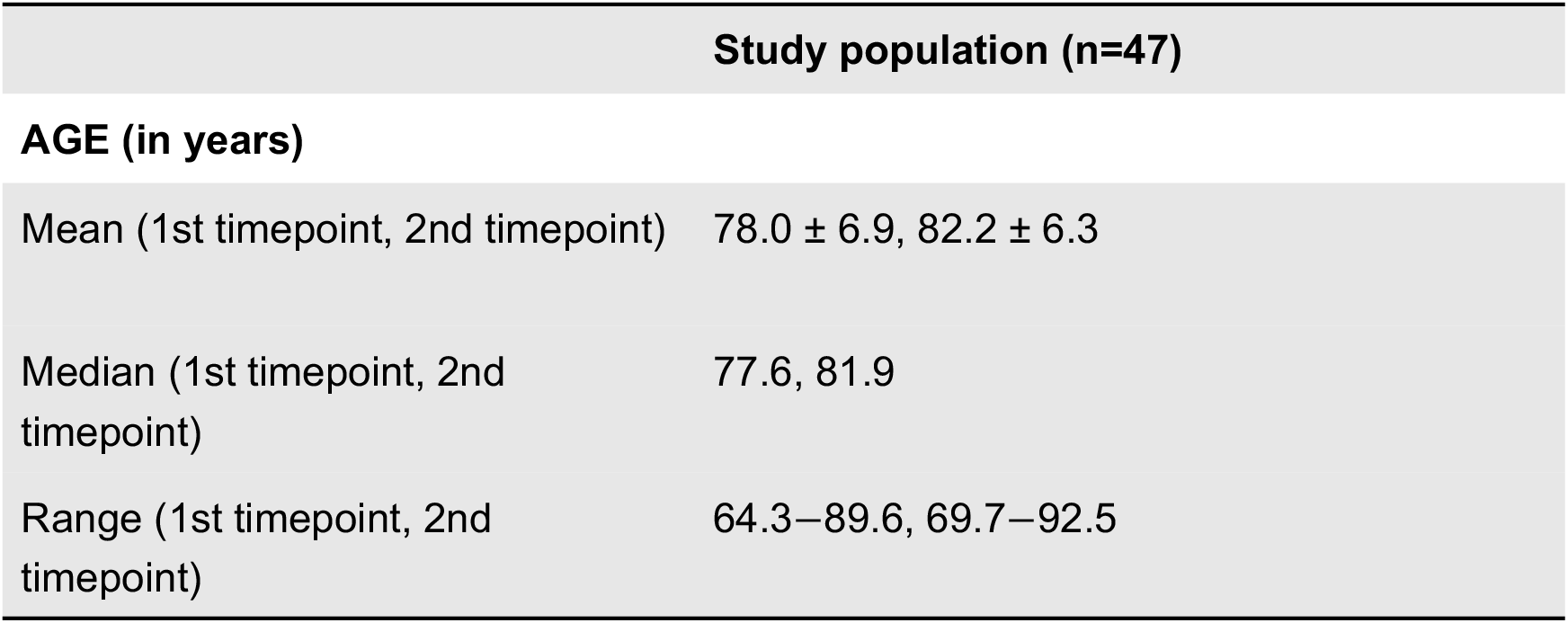
Demographic information for the BLSA Cohort. Table lists age at first and second timepoints.

| Study population (n=47) |  |
| --- | --- |
| AGE (in years) |  |
| Mean (1st timepoint, 2nd timepoint) | 78.0 ± 6.9, 82.2 ± 6.3 |
| Median (1st timepoint, 2nd timepoint) | 77.6, 81.9 |
| Range (1st timepoint, 2nd timepoint) | 64.3–89.6, 69.7–92.5 |

**Table S3.** Antibody responses against specific ToxScan peptides associate with the MHC class-II locus on chromosome 6 and NXN on chromosome 17. GWAS discovery was performed on individuals of European descent (EUR) in the VRC cohort. The findings were replicated in 3 of the 4 peptides in individuals of African descent (AFR) with p-values provided for both cohorts.

| Organism | Protein | Peptide position | Associated gene | SNP | Chr | Position | Ref/ Alt | OR EUR (95% CI) | EUR p-val | OR AFR (95% CI) | AFR p-val |
| --- | --- | --- | --- | --- | --- | --- | --- | --- | --- | --- | --- |
| *Staphylococcus aureus* | Clumping factor A (ClfA) | 505-560 | HLA-DQA1,HLA-DQB1 | rs539981616 | 6 | 32619171 | C/G | 6.10 (3.92 - 9.49) | 1.1E-15 | 23.42 (4.88 - 112.43) | 8.2E-05 |
| *Staphylococcus aureus* | Staphylococcal secretory antigen (SsaA1) | 1-56 | HLA-DQB1 | rs111951747 | 6 | 32627242 | C/T | 7.32 (3.94 - 13.59) | 2.8E-10 | 106.16 (6.98 - 1614.43) | 7.8E-04 |
| *Staphylococcus aureus* | Clumping factor A (ClfA) | 533-588 | HLA-DQA1,HLA-DQB1 | rs539981616 | 6 | 32619171 | C/G | 6.10 (4.00 - 9.32) | 5.8E-17 | 25.02 (4.93 - 127.06) | 1.0E-04 |
| *Streptococcus pyogenes* serotype M3 | Streptolysin O (SLO) | 85-140 | NXN | rs2663319 | 17 | 790803 | G/A | 0.05 (0.02 - 0.13) | 2.4E-10 | 3.71 (0.07 - 193.77) | 0.52 |

**Table S4.** Demographic information for the Crohn’s cohort. Table lists sex, age, Harvey-Bradshaw index of Crohn’s disease severity, and the Montreal classification of Crohn’s disease location.

| Study population (n=50) |  |
| --- | --- |
| **SEX** |  |
| Male | 19 (38%) |
| Female | 31 (62%) |
| **AGE (in years)** |  |
| Mean | 41.3 ± 14.5 |
| Median | 42 |
| Range | 18–75 |
| **HARVEY-BRADSHAW INDEX** |  |
| Mean | 10.5 ± 4.5 |
| Median | 9 |
| Range | 4–21 |
| **CROHN'S LOCATION** |  |
| L1: Ileal | 25 (50%) |
| L3: Ileocolonic | 25 (50%) |

**Table S5.** Demographic information for the JDM cohorts. Table lists sex, age at time of sampling, and race.

JDM
| Study Population (n=23) |  |
| --- | --- |
| **SEX** |  |
| Male | 8 (34.8%) |
| Female | 15 (65.2%) |
| **AGE (in years)** |  |
| Mean | 8.5 ± 3.5 |
| Median | 8 |
| Range | 3–17 |
| **RACE** |  |
| Black | 1 (4.3%) |
| Hispanic or Latino | 2 (8.7%) |
| Multiracial | 1 (4.3%) |
| White | 19 (82.6%) |

JDM Age-Matched Controls
| Study Population (n=23) |  |
| --- | --- |
| **SEX** |  |
| Male | 8 (34.8%) |
| Female | 15 (65.2%) |
| **AGE (in years)** |  |
| Mean | 9 ± 4 |
| Median | 9 |
| Range | 3–19 |
| **RACE** |  |
| Black | 1 (4.3%) |
| Hispanic or Latino | 1 (4.3%) |
| Multiracial | 1 (4.3%) |
| White | 19 (82.6%) |
| Unknown | 1 (4.3%) |

**Table S6.** Demographic information for the IBM cohorts. Table lists sex, age at time of sampling, and race.

IBM1
| Study Population (n=148) |  |
| --- | --- |
| **SEX** |  |
| Male | 90 (60.8%) |
| Female | 58 (39.2%) |
| **AGE (in years)** |  |
| Mean | 67.5 ± 8.0 |
| Median | 68 |
| Range | 43–89 |
| **RACE** |  |
| Asian | 1 (0.7%) |
| Black | 8 (5.4%) |
| Other | 2 (1.3%) |
| White | 125 (84.5%) |
| Unknown | 12 (8.1%) |

IBM2
| Study Population (n=57) |  |
| --- | --- |
| **SEX** |  |
| Male | 30 (52.6%) |
| Female | 27 (47.4%) |
| **AGE (in years)** |  |
| Mean | 60 ± 9 |
| Median | 60.5 |
| Range | 32.0–79.5 |
| **RACE** |  |
| Black | 11 (19.3%) |
| Hispanic or Latino | 11 (19.3%) |
| Multiracial | 35 (61.4%) |

